# Transposable elements drive phenotypic variation and shape the response to environmental changes in *Drosophila melanogaster*

**DOI:** 10.64898/2026.08.24.746693

**Authors:** Anaïs Larue, Alexander Mauro, Miriam Merenciano, Sonia Janillon, Florian Blanchard, Agnès Vallier, Alfredo Escanciano-Gómez, Marie Fackeure, Sandrine Hughes, Patricia Gibert, Cameron K. Ghalambor, Séverine Chambeyron, Rita Rebollo, Cristina Vieira

## Abstract

Transposable elements (TEs) are ubiquitous repetitive DNA sequences that can mobilise within genomes and may modulate gene expression in an environment-dependent manner. TEs and the safeguarding epigenetic machinery targeting them, can be tuned by environmental fluctuations to influence gene expression by inducing genomic, epigenetic, and transcriptomic changes. Yet, the degree to which TE-driven molecular diversity translate into inter-individual phenotypic variation *vs* accumulating without any phenotypic consequences remains unclear. Here, we used five populations of genetically engineered *Drosophila melanogaster* flies that carry variable TE content but share an otherwise identical genetic background to test the phenotypic consequences of the early stages of TE accumulation. Phenotypic screenings across 17 traits (fertility-related traits, life-history traits and stress resistance tests) revealed significant differences between the populations (e.g. reduced hatchability). We also observed a notable increase in intra-population phenotypic variation for the heavily TE-burdened populations across a wide panel of traits. These results suggest considerable TE-driven inter- and intra-population phenotypic variation. Further investigation revealed that variable TE contents can influence the response to environmental changes, positioning TEs as drivers of environmentally-induced phenotypic variation in a system deprived of other sources of genetic variation. These results provide empirical evidence that TEs contribute to the heterogeneity of the environmental response and therefore represent an underlying mechanism of phenotypic variation.

**Significant statement:** The molecular mechanisms underlying phenotypic variation and variation in the environmental response continue to be a central question in evolutionary biology. Leveraging a biological system composed of *Drosophila melanogaster* populations containing varying levels of transposable elements (TEs), which are repetitive and widespread genomic elements, within an otherwise identical genetic background, we demonstrate the influence of TEs in generating phenotypic variation. By employing this innovative system to isolate the specific contributions of TEs, we provide evidence for their role as drivers of both phenotypic variability and divergence of the environmental response. Such results show that, rather than being largely neutral or silenced through epigenetic mechanisms, TE accumulation can contribute to measurable phenotypic diversity and thus provide heritable variation that selection can act upon.

## Introduction

Transposable elements (TEs) are ubiquitous repetitive DNA sequences capable of mobilising within most eukaryotic genomes (Wells and Feschotte 2020). Although TEs are most often neutral or deleterious, they have in some cases contributed to the emergence of adaptive phenotypes (Rey et al. 2016; Platt et al. 2018; Schrader and Schmitz 2019; Catlin and Josephs 2022; Galbraith and Hayward 2023). The potential of TEs to drive adaptation by generating genetic and phenotypic variation stems from two key properties. First, TEs can contribute to phenotypic diversity by carrying regulatory elements capable of influencing the expression of nearby genes, acting as promoters or enhancers, or supplying transcription factor binding sites (TFBSs) (Chuong et al. 2017; Dubin et al. 2018; Drongitis et al. 2019; Moschetti et al. 2020; Sundaram and Wysocka 2020; Gebrie 2023). TE insertions can also generate new polyadenylation signals, introduce premature stop codons, or give rise to chimeric transcripts (Oliveira et al. 2023; Huang and Lee 2024). Secondly, because TEs are silenced by epigenetic mechanisms to counteract their deleterious effects, they can contribute to cryptic variation through epigenetic changes in DNA methylation, histone tail modifications, and small RNAs (sRNAs), that suppress nearby gene expression (Rebollo et al. 2012; Rey et al. 2016; Fueyo et al. 2022). Indeed, TE silencing mechanisms are known to influence the expression of nearby genes, a phenomenon documented in the literature as epigenetic spreading (Guio et al. 2018; Choi and Lee 2020; Huang et al. 2022).

These TE-driven transcriptomic, epigenetic, and phenotypic impacts can also be environmentally-induced. For example, in *D. melanogaster*, the insertion of a solo-LTR of a *roo* element upstream of the *Lime* transcription factor (TF) gene was reported to introduce a functional alternative promoter to *Lime* and to impact the responses to immune stress (Merenciano and González 2023). Upon bacterial infection with *Pseudomonas entomophila, D. melanogaster* flies carrying the *roo* insertion experience an increased survival, driven by the upregulation of the *Lime* gene mediated through *roo*’s dual role as promoter and enhancer by the presence in its sequence of TFBSs (Merenciano and González 2023).

Environmental changes can modulate TEs’ activity and their epigenetic control, which may influence the response of the organism to environmental variation. As a result, individuals with variable TE composition may exhibit distinct TE-driven phenotypes depending on the environment. While some studies have linked discrete TE insertions to specific phenotypic changes (Schrader and Schmitz 2019; Gilbert et al. 2021), the contribution of an organism’s overall TE content remains largely unexplored. Moreover, it is impossible to distinguish, in a wild-type system, the contribution of TE to the phenotype from the effects of genetic background variations. This reflects our ability to detect the end-point evolutionary contributions of TEs to phenotypes, while also highlighting the difficulty in evaluating their contribution at shorter evolutionary timescales. Consequently, a number of conceptual papers have emerged theorising on the contribution of TEs to environmentally-driven phenotypic variation (Piacentini et al. 2014; Lanciano and Mirouze 2018; Marin et al. 2020; Pimpinelli and Piacentini 2020), and hence to generating raw material for selection, with little to no empirical evidence to support it.

To address this gap, we designed an experimental approach in which sources of genetic variation other than TE insertions are minimized, to isolate their specific contribution to the phenotype. Within this framework, we tested whether TE content and its variation across populations could account for part of the observed phenotypic divergence, both between populations (inter-population variation) and within a single population (intra-population variation). We also assessed to what extent differences in TE content and position could modulate the phenotypic response to environmental changes. To do that, we used a genetically modified isogenic *D. melanogaster* line (G0), from which four populations were derived (G10, G31, G73, and G100). Each of the populations is the result of a varying number of generations of inducible somatic *Piwi* knockdown (KD) (Barckmann et al. 2018; Varoqui et al. 2025). Indeed, the *Piwi* protein plays a central role in the *piwi*-interacting RNA (piRNA) pathway that regulates transcriptional TE activity in *Drosophila*, and its successive KD resulted in TE accumulation, all within an otherwise constant genetic background (Barckmann et al. 2018; Varoqui et al. 2025). Within this framework, G10 corresponds to ten successive generations of inducible *piwi* KD in a population of around 500 individuals, resulting in polymorphic TE accumulation, while G31 corresponds to 31 generations, and so on. This provided us with a controlled experimental framework in which the resulting TE-driven polymorphism could be leveraged to evaluate the contribution of TEs to phenotypic variation and environmental response across populations.

Here, we tested the impact of differences in TE content on a panel of phenotypic traits, including fitness indicators (hatchability and viability), behavioural assays (climbing), and life history traits (longevity and developmental time). The phenotypic screening revealed significant differences between populations, including a notable decrease in fitness in the G73 flies. An analysis of the variance across all traits, using the coefficient of variation (CV), revealed that the fitness reduction in G73 was accompanied by a general increase in phenotypic variability. This suggests that a greater intra-population insertional polymorphism results in dispersion of the phenotypes around the population mean, which is a requirement for an evolutionary response. Additionally, exposing the TE-divergent populations to diverse environmental conditions revealed that distinct TE contents, within an identical genetic background, resulted in distinctive environmental responses. Specifically, significant differences were observed in longevity under fluctuating diets and temperatures, as well as in stress tolerance across varying oxidative intensities. These findings support that TEs take part in shaping the environmental response. Finally, while overall TE content contributes to phenotypic differences between populations, we also identified candidate insertions that may provide the mechanistic basis for given phenotypic divergences, through a genome-wide scan approach. To our knowledge, this is the first study to provide evidence for the contribution of TEs to phenotypic variation under such a tightly controlled experimental framework in insects.

## Results and Discussion

### The TE-divergent populations differ in TE content and composition

A growing number of studies have implicated TEs in playing a key role in adaptive evolution, leading many to conclude that TEs are playing an underappreciated role in the evolutionary responses to natural selection (Casacuberta and González 2013; Schrader and Schmitz 2019; Marin et al. 2020; Gilbert et al. 2021; Merenciano et al. 2024). However, capturing the initial role TEs play in generating this heritable phenotypic variation has been challenging because of the unpredictability of TE activity across the genome and the lack of adapted tools. Here, we use an experimental approach that took advantage of an isogenic line of *D. melanogaster* in which different levels of TE accumulation have been induced to establish pools of flies with variable amounts of TE accumulation, so-called populations (see Materials and Methods). This enabled us to directly investigate the contribution of TEs to phenotypic variation under both control and stressing environmental conditions. (Barckmann et al. 2018; Mohamed et al. 2020; Varoqui et al. 2025). To accurately describe the overall TE content and composition of the TE-divergent populations, hereafter jointly referred to as TE composition, we employed the TrEMOLO pipeline (Mohamed et al. 2023). The OUTSIDER detection module from TrEMOLO enables the detection of polymorphic TEs and estimates their frequencies in the population by mapping long reads against an assembly (Mohamed et al. 2023). Using this module on high-coverage long read sequencing data (∼60 X to 140 X), Varoqui and collaborators compared the number of new insertions of Long Terminal Repeat (LTR) retrotransposons in G11 (corresponding to 11 generations of TE accumulation), G31 (31 generations), G73 (73 generations) relative to the G0 parental population (no TE accumulation). They detected 280, 514, and 798 novel LTR insertions from 43 families, in G11, G31, and G73 respectively (Varoqui et al. 2025). We have acquired a subset of these populations, in addition to a G100 population (100 generations of TE accumulation), and confirmed the accumulation of TE insertions across generations along with high insertional polymorphism by long read DNA sequencing (35 X coverage). Unlike previous descriptions focusing on LTR elements, we report insertions across all TE classes, including unexpected increases in LINE elements. From a panel of 165 consensus TE sequences described in *D. melanogaster* present in the most recent manually-curated TE library (Rech et al. 2022), TrEMOLO identified 373, 414, 496 and 518 new TE insertions in the G10, G31, G73 and G100 populations respectively, compared to the G0 (Figure 1A), confirming an increasing accumulation of new TE insertions across generations (complete analysis available in supplementary material: see Figures S1, S2 and S3). The newly inserted TEs belong to 79 different TE families, suggesting they are transpositionally active in the G0 parental line and may have a basal transposition rate. Considering all newly detected insertions, the average insertion frequency estimated with TrEMOLO (Mohamed et al. 2023) range from approximately 13% to 17% across the populations (Table 1), suggesting high TE insertion polymorphism in agreement with Varoqui (Varoqui et al. 2025). It is worth noting that global frequency distributions did not differ significantly between populations (Kruskal-Wallis p-value = 0.0659).

**Figure 1:**
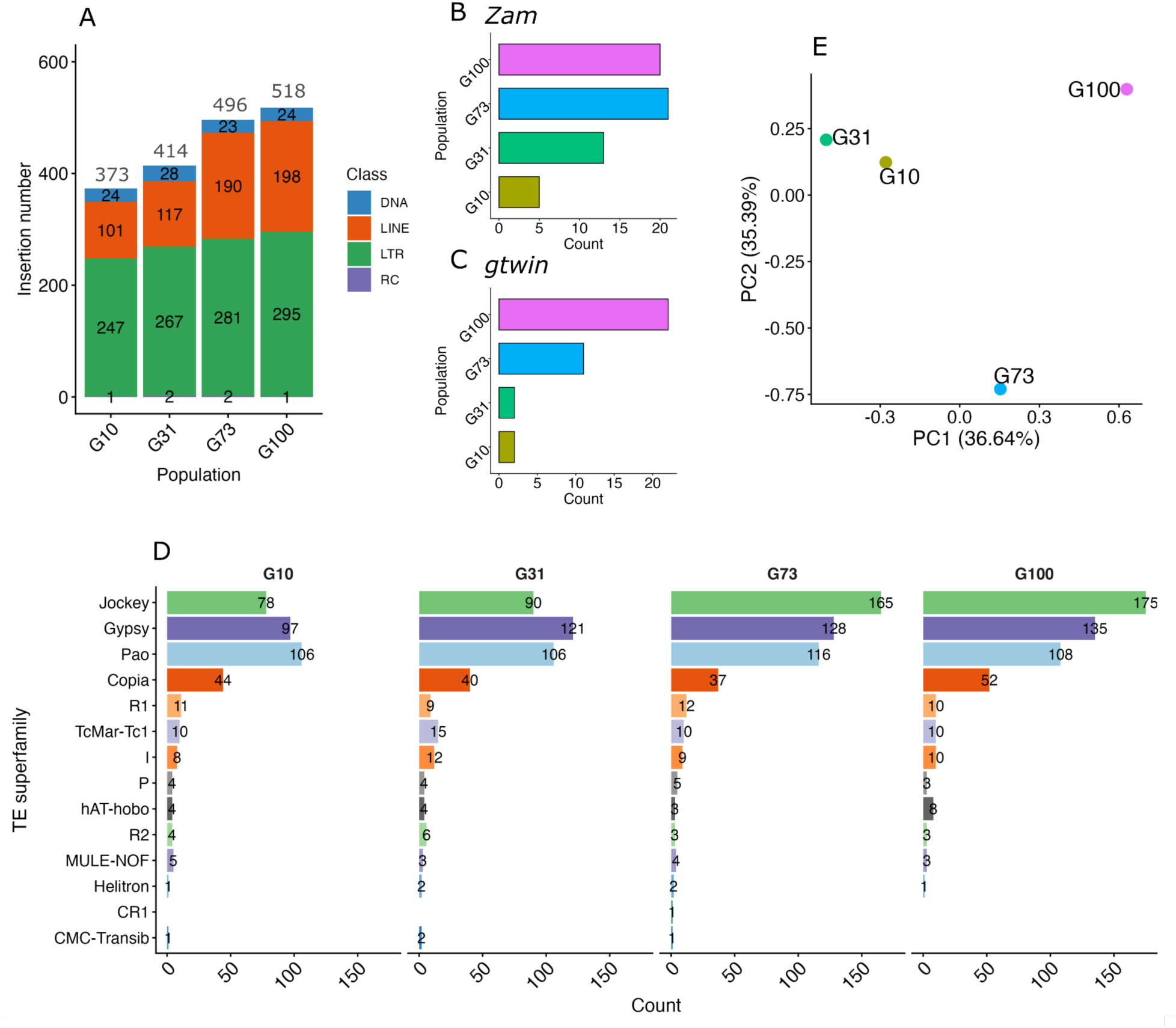
TE content increases in the TE accumulation populations. A) Distribution of new TE insertions across populations compared to G0 parental line detected by TrEMOLO. The number in each of the colour categories corresponds to the number of insertions per subclass. Distribution of *Zam* (B) and *gtwin* (C) new insertions in each population. E) New TE insertions at the family level. Numbers indicate the TE insertions detected by TrEMOLO. D) Principal component analysis of the TE frequencies.

**Table 1:**
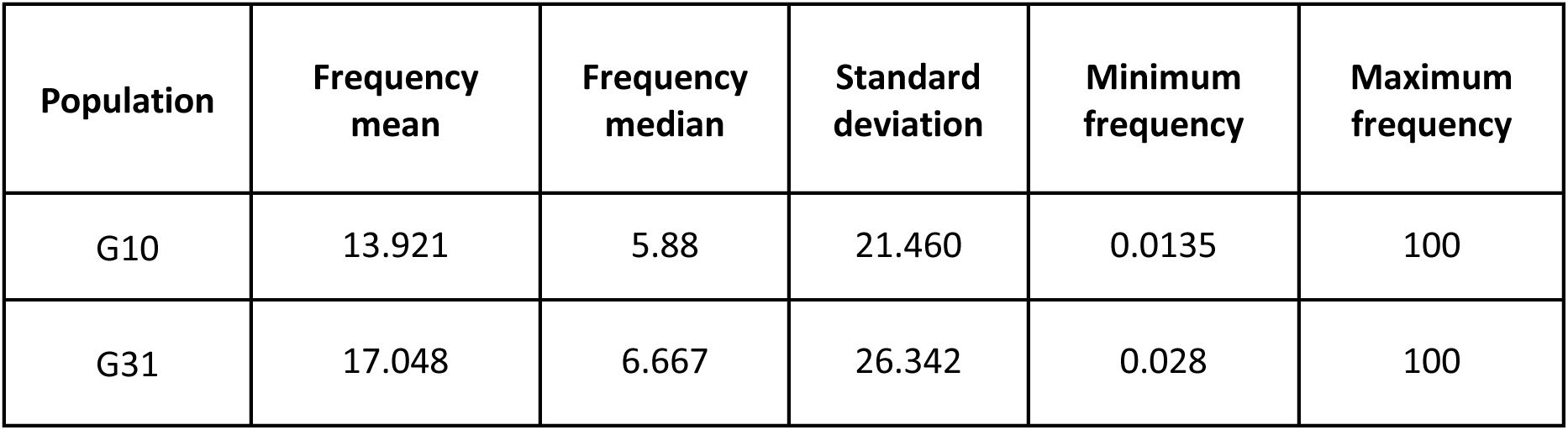
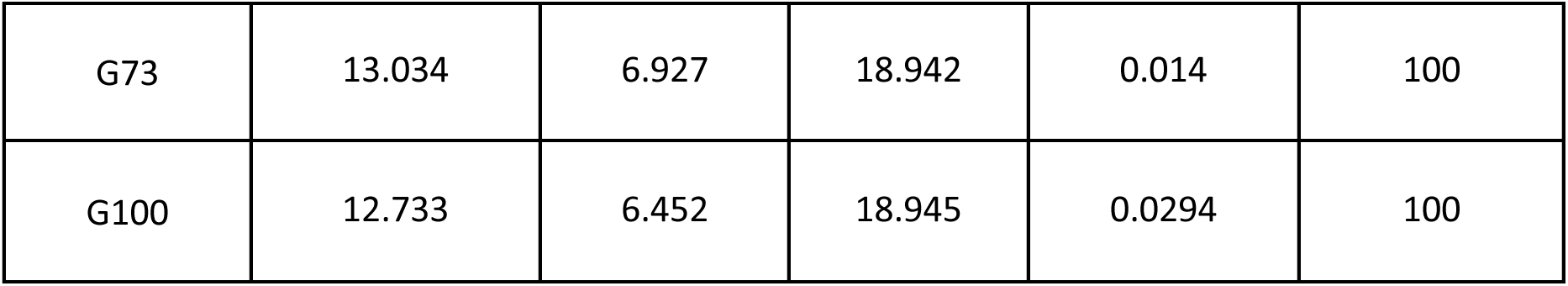
Summary statistics of the insertion frequencies at the population scale. Full-length TE insertions, not present in G0 parental line, were detected using TrEMOLO in G10, G31, G73, and G100 populations, which also inferred insertion frequencies for each population. The table reports the mean, median, standard deviation, and the minimum and maximum frequency of TE insertions for each population.

Regarding LTR elements, new insertions increase by 247, 267, 281 and 295 in populations G10, G31, G73 and G100, respectively (Figure 1A, Figure S1). Within this TE subclass, *Zam* and *gtwin* display significant increase in the number of new insertions compared to G0 parental genome, confirming previous results (Barckmann et al. 2018; Varoqui et al. 2025) (Figure 1A, 1B and 1C). Moreover, the TE-divergent populations also display an increase in LINE elements, (Figure 1A, Figure S2). LINE elements increased by 101 in G10, and by 117, 190 and 198 in G31, G73 and G100, respectively (Figure 1A). For instance, *Jockey* family (LINE element) continuously increases until generation 100 (G10=78, G31=90, G73=165 and G100=175) (Figure 1D). More specifically, two LINE elements within the *Jockey* family, namely *F-element* and *Doc,* display an important increase in new insertions across generations (Figure S2). As for DNA and Rolling Circle (RC) transposons, no noticeable increase was observed (Figure 1A, Figure S3).

Additionally, we used the insertion frequencies from TrEMOLO to infer the variation in TE composition of the four populations derived from the G0 parental line. We first performed a principal component analysis (PCA) on the insertion frequencies (Figure 1E), separating the populations into three groups. PC1 explains 36.64% of the variance and clusters G10 and G31 together, followed by G73 and G100 (Figure 1E) while PC2 explains 25.43% of the variance and separates G73 from the other populations (Figure 1E), highlighting the insertional polymorphism between populations, and adding a layer of variability beyond mere TE content accounting for TE insertion frequency.

Finally, an analysis of non-repeat structural variants (SVs) (non-TE and non-satellites SVs) was conducted to evaluate the extent to which they could contribute to the genotypic variation across the TE-divergent populations (complete analysis available in supplementary material: see Supporting Information, Table S1 and Figures S4, S5). With an average of 114 indels between the TE-divergent populations relative to G0 (Table S1), the numbers of non-repeat SVs detected between the genome assemblies are at least one order of magnitude below those reported in the literature when comparing *D. melanogaster* assemblies (Chakraborty et al. 2018, 2019; Liu et al. 2025). For further insight, a telomere-to-telomere comparison of ISO-1 and Canton S assemblies, detected 7,989 SVs (Liu et al. 2025). Notably, only 1% of the detected SVs belonged to repetitive sequences (Liu et al. 2025). This reveals that TE composition is the parameter showing the most considerable variation between the present populations.

Taken together, the variation in TE composition, along with their potential regulatory and structural implications, makes these populations an ideal experimental system for evaluating the impact of TEs on phenotypic variation under both controlled and changing environmental conditions.

### Variable TE content influence phenotypic profiles across populations

Having confirmed the TE accumulation pattern across the populations as well as demonstrated their distinct TE compositions, all within an otherwise nearly identical genetic background, we aimed to investigate their phenotypic impact. Given that TE composition is the main genetic variation between the populations, we expect that any phenotypic divergence will be driven by TEs, either by their global or insertion-specific effects. Thus, a broad range of traits were measured, aimed at examining various biological functions. These included traits directly contributing to fitness, such as hatchability, viability, and developmental time, as well as life-history traits like longevity and dry weight, all of which showed differences between the TE-divergent populations, statistically supported by a strong population effect (Table 2). Additionally, metabolic rate was measured to provide a broader view of the physiological impact of TEs, alongside a behavioural assay (climbing ability), and survival under oxidative and thermal stress. The metabolic rate measurements revealed population divergence as temperature increased both in females and males (Table 2). Climbing abilities were only slightly different for the males (Table 2). Finally, oxidative stress and thermal changes also highlighted significant population effects on the phenotypic outcome (Table 2). Briefly, out of the 17 traits measured, a population effect, *i.e.* the effect of TE composition, was observed in six female traits, seven male traits and two traits with undetermined sex (mixed) (all p-values < 0.001) (Table 2). Some examples of these phenotypic divergences are discussed in further detail below, but three major findings emerged: G73 showed the strongest fitness costs (reduced hatchability and viability, p < 0.001), G31 had delayed development (21.2 *vs* 20 days, p < 0.001) but an increased longevity at 18°C for the females (p < 0.001), and G10 males were smaller (dry weight, p < 0.01), correlating with reduced lifespan.

**Table 2:**
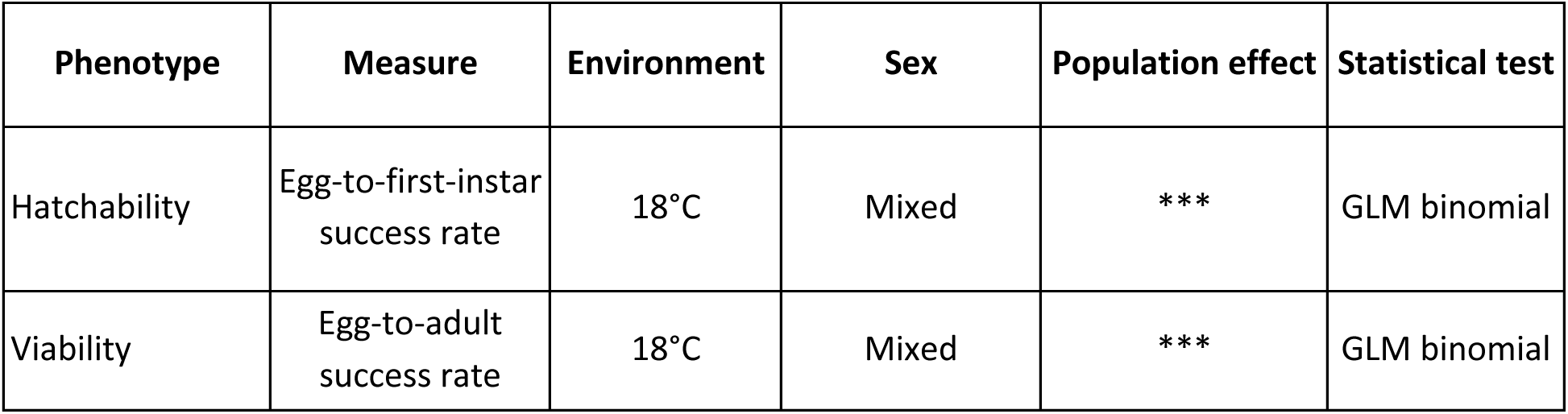
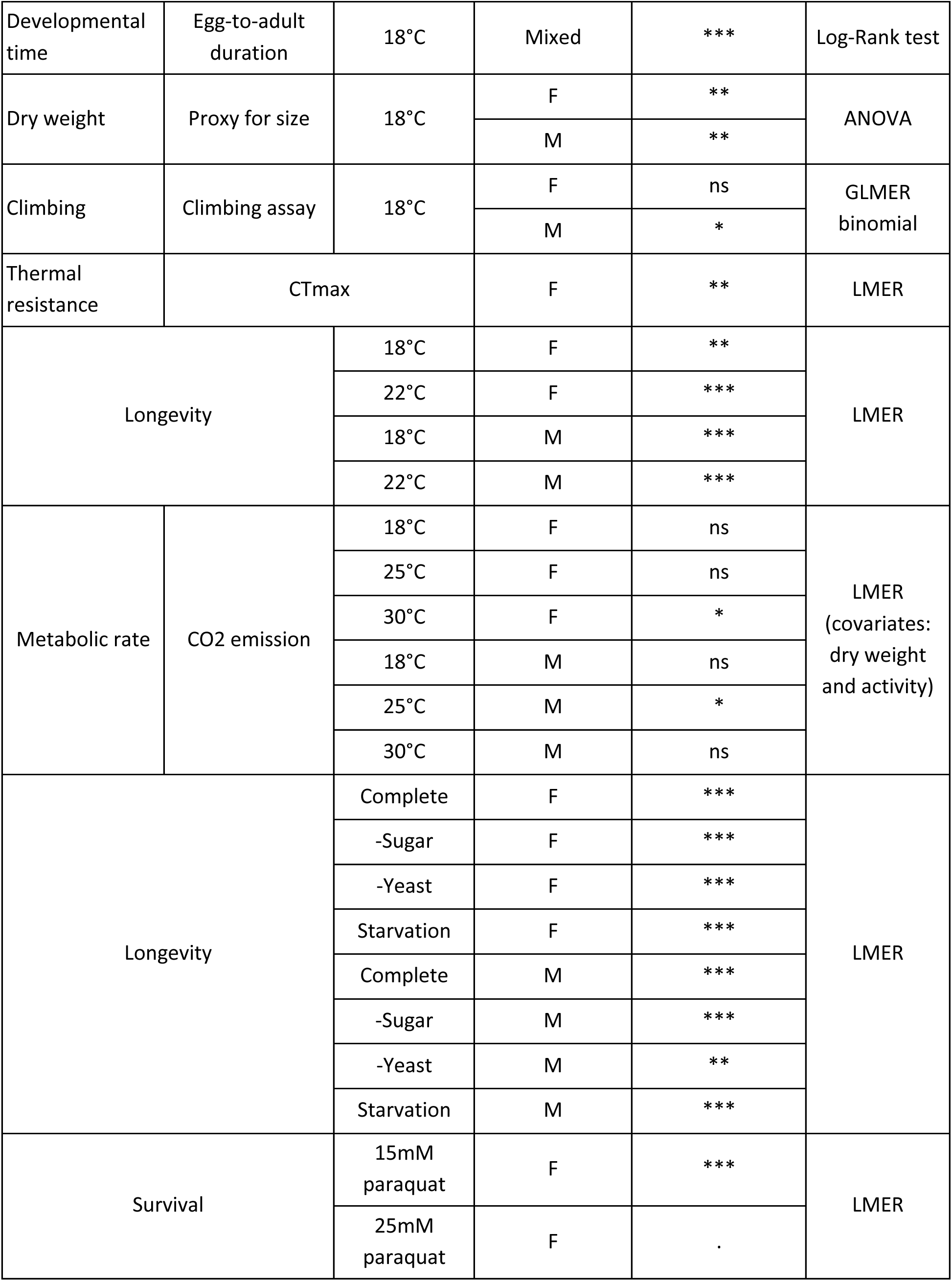
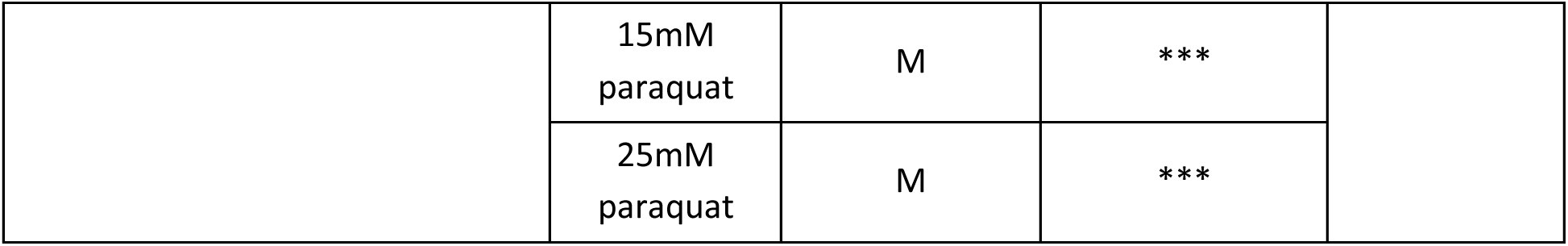
Summary of phenotypic measurements across populations, sexes, and environmental conditions. The “Phenotype” column indicates the trait assessed, “Measure” specifies how the trait was quantified, and “Environment” indicates the conditions under which the measurement was performed. “Sex” identifies whether the measurement was made on females (F), males (M), or mixed. “Population effect” denotes the statistical significance of differences among populations. “Statistical test” specifies the model used to assess the population effect, including generalized linear models (GLM), linear mixed-effects models (LMER), generalized linear mixed-effects models (GLMER), and Log-Rank tests for time-to-event data. For the metabolic rate, the LMER analyses accounted for dry weight and activity as covariates. - Sugar and -Yeast designate the nutritional source depleted from the media. Statistical significance is depicted as follows: 0 ‘***’ 0.001 ‘**’ 0.01 ‘*’ 0.05 ‘.’ 0.1 ‘ns’ 1

Among the most striking differences was the significant decrease in hatchability (the proportion of eggs from which a larva emerges) in the G31 and G73 compared to the G0 and G10 (GLM p-values = 0.0002, 0.0032, respectively) (Figure 2A). The decrease in hatchability observed is in agreement with the findings of Pasyukova et al. (2004), who reported a decreased hatchability in a *D. melanogaster* strain carrying up to 90 additional TE insertions of the *copia*, *roo,* and *P-element* families compared to other lines. They propose that the effect of TEs on hatchability may be a consequence of higher rates of deleterious chromosomal rearrangements and in an increased embryonic death (Pasyukova et al. 2004).

**Figure 2:**
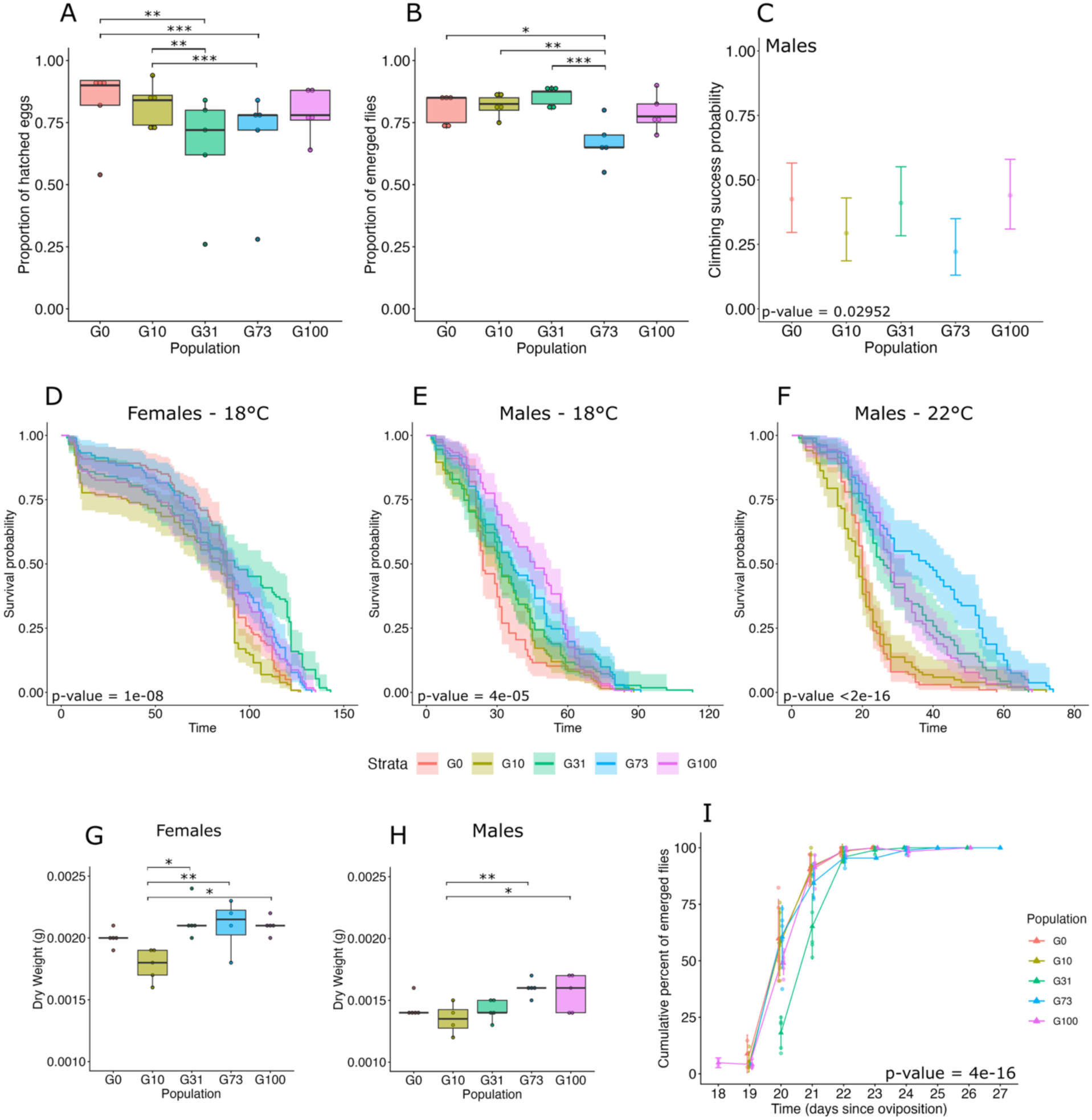
Examples of phenotypic divergences between the TE-divergent populations. A) Hatchability expressed in proportion of hatched eggs into larvae. Each condition is composed of five replicates of 50 eggs. G31 and G73’ hatchability appears reduced compared to both G0 and G10. B) Viability, representing the proportion of survival from egg to adult. Each condition consisted of five replicates of 40 initial eggs. C) The fitted climbing success probability for the flies of each population to have reached the top of the tube at the end of the trial. Each condition was replicated six times with pools of 10 flies. D)-F) Longevity measured as the survival curves of the females at 18°C (D) and the males at 18°C (E) and at 22°C (F). The dry weight of the females (G) and males (H). Each of the conditions is represented by five replicates of a pool of five flies. I) Cumulative percent of emergent flies as a function of the day since oviposition. Each condition comprised five replicates of 40 initial eggs.

In addition, a decrease in viability (measure of the egg-to-adult survival) in the G73 compared to all other populations suggest that not only hatchability is impacted by TE load, but later developmental stages are also affected in this population (Figure 2B). Viability in optimal condition is expected to range from 70% to 80% in *D. melanogaster* (Flatt 2020). Coherent with our observation, previous fitness measurements conducted on three lines with varying *P-element* content revealed that viability decreased in strains increasing with *P-elements* insertions (Mackay 1986).

Finally, the climbing assays show a significant population effect (GLMER p-value = 0.0295) with marginally significant lower propensity to reach the upper part of the test vial for the males G73 compared to the males G100 (GLMER p-value = 0.0556) (Figure 2C). No significant differences were reported for the females. It was previously reported that males with somatically active *P-elements* have shown lower locomotion than their *P-element*-free counterparts (Woodruff et al. 2000). We hypothesized that the marked increase in TE content of +82 insertions detected from G31 to G73 (414 and 496 insertions, respectively), followed by a plateau at G100 (+22 insertions compared to G73, with 518 TEs detected), could reflect a specific turnover in TE insertions at G73, acting as a burden and potentially manifesting as an impairment in males’ locomotion. Thus, we find evidence that TE load has some deleterious fitness consequences on some traits.

While the three previously described phenotypes appear to reflect the influence of overall TE load, without showing a linear correlation with its increase *per se*, the subsequent phenotypes seem to be driven by population-specific insertions, highlighting the potential targeted effects of particular TE insertions on phenotypic differences between populations. First, longevity experiments show no general pattern of correlation to the increase in TE content. For example, at 18°C G10 females experience a decrease in longevity compared to G31, G73 and G100 females (Log-Rank test p-values = 7.7e-07, 0.0031 and 0.018, respectively) (Figure 2D). The longevities of G0 and G10 males at 18 °C are also reduced compared to those of G73 and G100 (Log-Rank test p-values: 0.0397 and 0.0027 for G0 *vs.* G73 and G100, respectively; 0.0048 and 0.0004 for G10 *vs.* G73 and G100, respectively) (Figure 2E). The decrease in lifespan of G0 and G10 males appears to be emphasized at 22°C, while G73’s is drastically increased (Figure 2F).

The dry weight appears as a potential predictor of the G10’s reduced lifespan (Figure 2G and 2H). In *D. melanogaster*, the relationship between body size and longevity is highly variable, however a positive correlation has been previously described for males of a wide panel of tested strains (Khazaeli et al. 2005). And indeed, G10 males are generally smaller (the dry weight is used as a proxy for size) than both G73 and G100 (ANOVA p-values = 0.0074 and 0.0202, respectively) (Figure 2H). Similarly, G10 females appear to be smaller than G31, G73 and G100 (ANOVA p-values = 0.0184, 0.0052 and 0.0418, respectively) (Figure 2G).

It is also worth noting the delay in development time of the G31 from the other populations (Figure 2I). Longer development time is often correlated with longer lifespan (Lints and Lints 1971; Mayer and Baker 1985; Stearns et al. 2000). Therefore, it is not unexpected to also observe an increase in longevity in the females of the G31 population compared to G0, G10 and G100 at 18°C (Log-Rank test p-values = 0.0005, 7.7e-07 and 0.0261, respectively) (Figure 2D). Indeed, the average emergence time for G31 is 21.23 days, while it lays around 20 days for the other populations as expected at 18°C (Pulver and Berni 2012) (Figure 2I). Therefore, in our experiments, G31 consistently takes one more day to reach adulthood. The difference in development time being specifically bound to one population may suggest that this is the result of one or more isolated TE variants’ effect, rather than related to the global TE content.

A principal component analysis (PCA) and pairwise Pearson’s correlation coefficients were also respectively computed to investigate the phenotypic spreading of the populations and the correlations between traits. In females, G0, G10 and G100 cluster together while G31 and G73 separate along PC1 and PC2 respectively (Figure 3A). Pearson’s coefficients reveal six significant positive correlations between phenotypes as well as six negative correlations (Figure 3B). Notably, development time seems to be positively correlated with longevity at 18°C (0.96), measured as the LT80 (the estimated time for 80% of the individuals to have died (see Material and Methods), as previously discussed (Figure 3B). In males, PCA analysis shows a slight clustering of G0 and G10 on one side, and of G73 and G100 and the other side, leaving G31 separated along PC2 (Figure 3C). Pearson’s coefficients reveal six positive correlations and one negative (Figure 3D).

**Figure 3:**
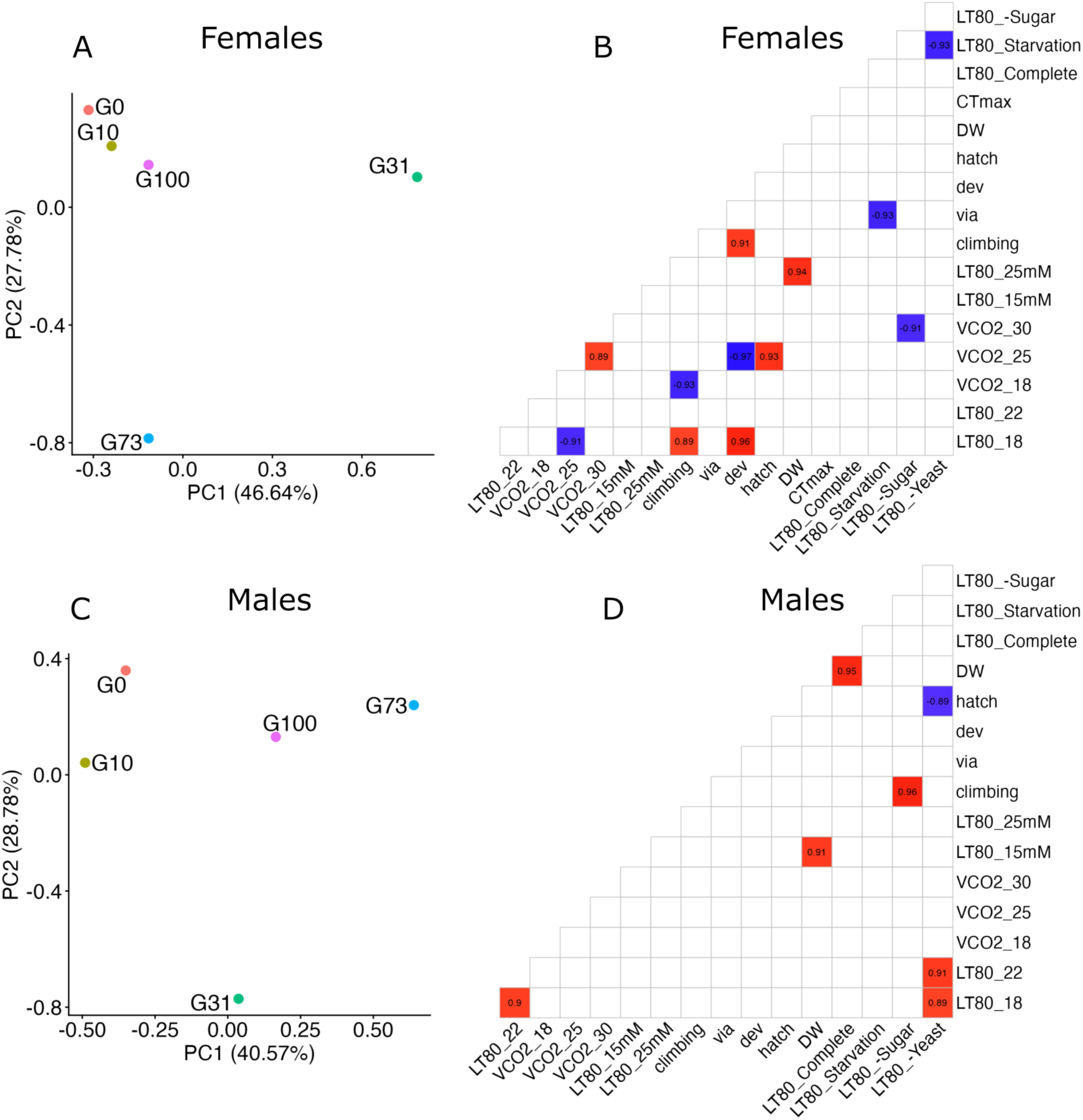
Dimensional and correlative data analysis of the phenotypes of the population with increasing TE content. A) PCAs of the means and LT80 values of all the phenotypic measurements and B) Pearson’s correlation matrixes of the p-value supported correlations for the means and LT80 values of all the phenotypic measurements for the females. Only the significant correlations are displayed. Similarly, C) PCAs of the means and LT80 values and D) Pearson’s correlation matrixes for the males. ”VCO2” refers to the metabolic rate measurements, “hatch”, to the hatchability, “via”, to the viability, “dev”, to the development time, “climbing”, to the climbing assay, “15mM” and “25mM” apply to the concentrations of paraquat, “DW” is the dry weight, “18”, “22”, “25”, “30” are the environmental temperature, and “Complete”, “Starvation”, “-Yeast”, “-Sugar”, the environmental media.

Additionally, for each phenotype, partial Eta^2^ and R^2^ metrics were calculated to estimate the proportion of variance explained by the TE composition (according to the statistical model) (Figure 4). Three phenotypes for males (longevity under complete and yeast deprivation diets, and locomotion) and five for females (both paraquat concentrations and starvation survival assays, longevity under both 18°C and 22°C and locomotion) fall below the threshold of mild effect (Figure 4). In females, four phenotypes are mildly to strongly impacted by the TE content (longevity under complete, sugar and yeast deprivation diets, and CTmax), while four for males (both paraquat concentrations and starvation survival assays and longevity at 18°C) and one for mixed (development time) (Figure 4). Altogether, 12 phenotypes are strongly impacted by the TE composition (both females’ and males’ metabolic rates under every temperature condition, both sexes’ dry weight, males’ longevity at 18°C and 22°C and under sugar deprivation diet, as well as hatchability and viability) (Figure 4).

**Figure 4:**
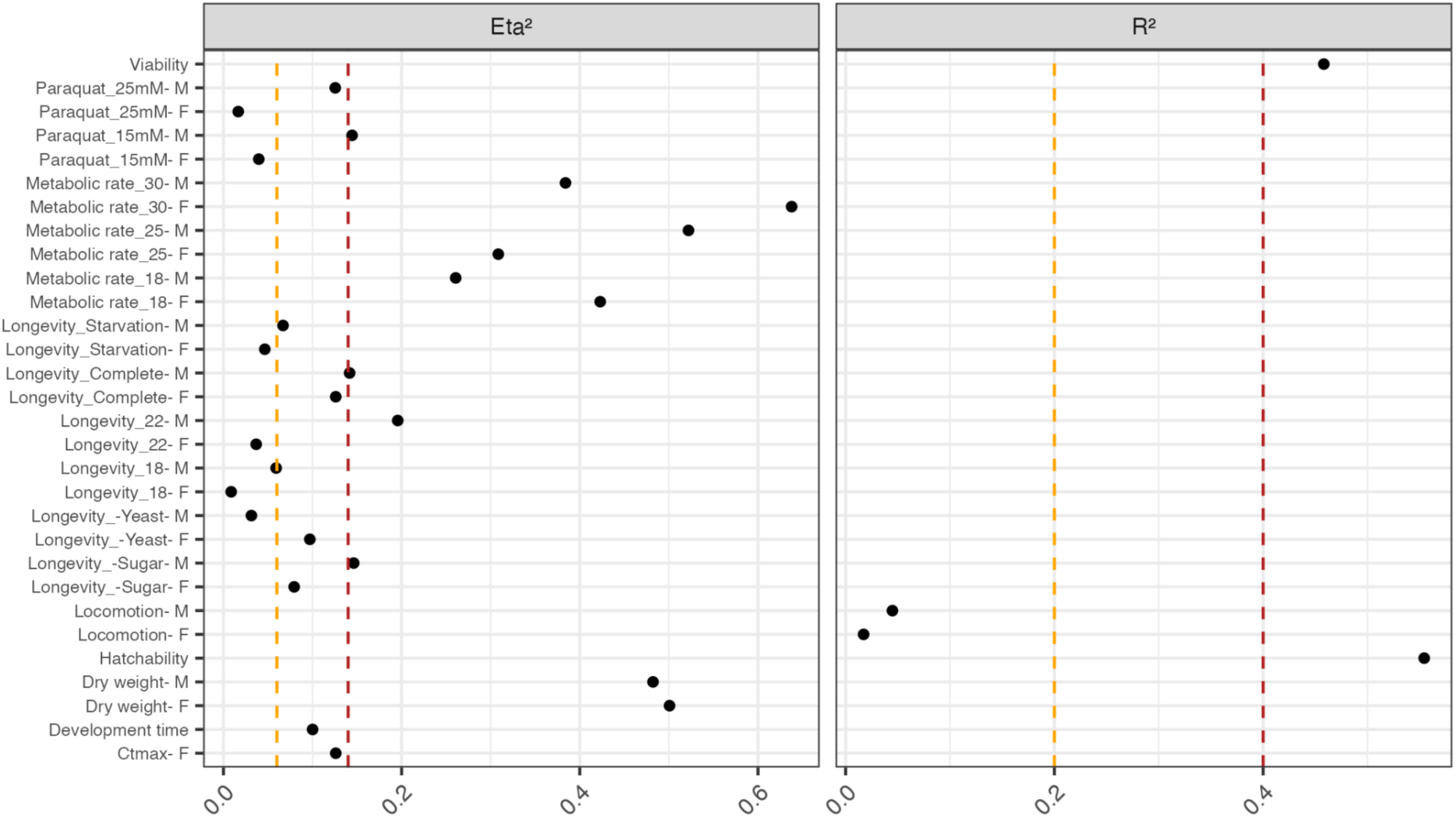
Summary of the effect sizes of the contribution of the population variable to the data. The contribution of the population variable to the observed data are expressed as the Eta² and the R² according to the nature of the data and statistical tests. The yellow vertical dashed line represents the threshold of medium effect and the red the threshold of strong effect. “F” refers to measures in females and “M” in males, “15mM” and “25mM” are the concentrations of paraquat, “DW”, the dry weight, “18”, “22”, “25”, “30” are the environmental temperature, and “Complete”, “Starvation”, “-Yeast”, “-Sugar”, the environmental media.

In conclusion, these results suggest that differences in TE composition in a nearly identical genetic background influences diverse phenotypes, highlighting the importance of considering the cumulative TE effects in *D. melanogaster.* However, the phenotypic measurements reveal that the impact of TEs is not as straightforward as a simple correlation between higher TE content and reduced performance. Instead, some of the significant differences observed across populations appear to reflect population-specific effects rather than a general correlation with TE abundance. These findings suggest that phenotypic variation may not solely arise from total TE content, but also from the presence of specific insertions contributing to phenotypic differentiation.

### High TE content increases the observed phenotypic variability in females

While the previous section shows that variation in TE composition drives inter-population phenotypic variation, we also investigated whether populations with higher TE content, and in this case, greater within-population insertional polymorphism, exhibit higher intra-population phenotypic variation. To test this hypothesis, the standard deviations (SD) from all phenotypic assays (used as a proxy for phenotypic variation to measure dispersion from the mean) were compiled (see Materials and Methods). A Friedman’s rank test was performed on the ranked SD of the females and showed that G73 SD were consistently greater than that of G0 and G10 (Friedman Rank test p-values = 0.020 and 0.017, respectively) (Figure 5A, Table S2). A similar analysis was performed on the male data, however, as it yielded no further statistical support, the subsequent analysis focused exclusively on the females. We computed the coefficient of variation (CV) for all the phenotypes, a measure of dispersion that expresses SD as a percentage of the mean and hence frees itself from the relationship to the mean (Morgante et al. 2015).

**Figure 5:**
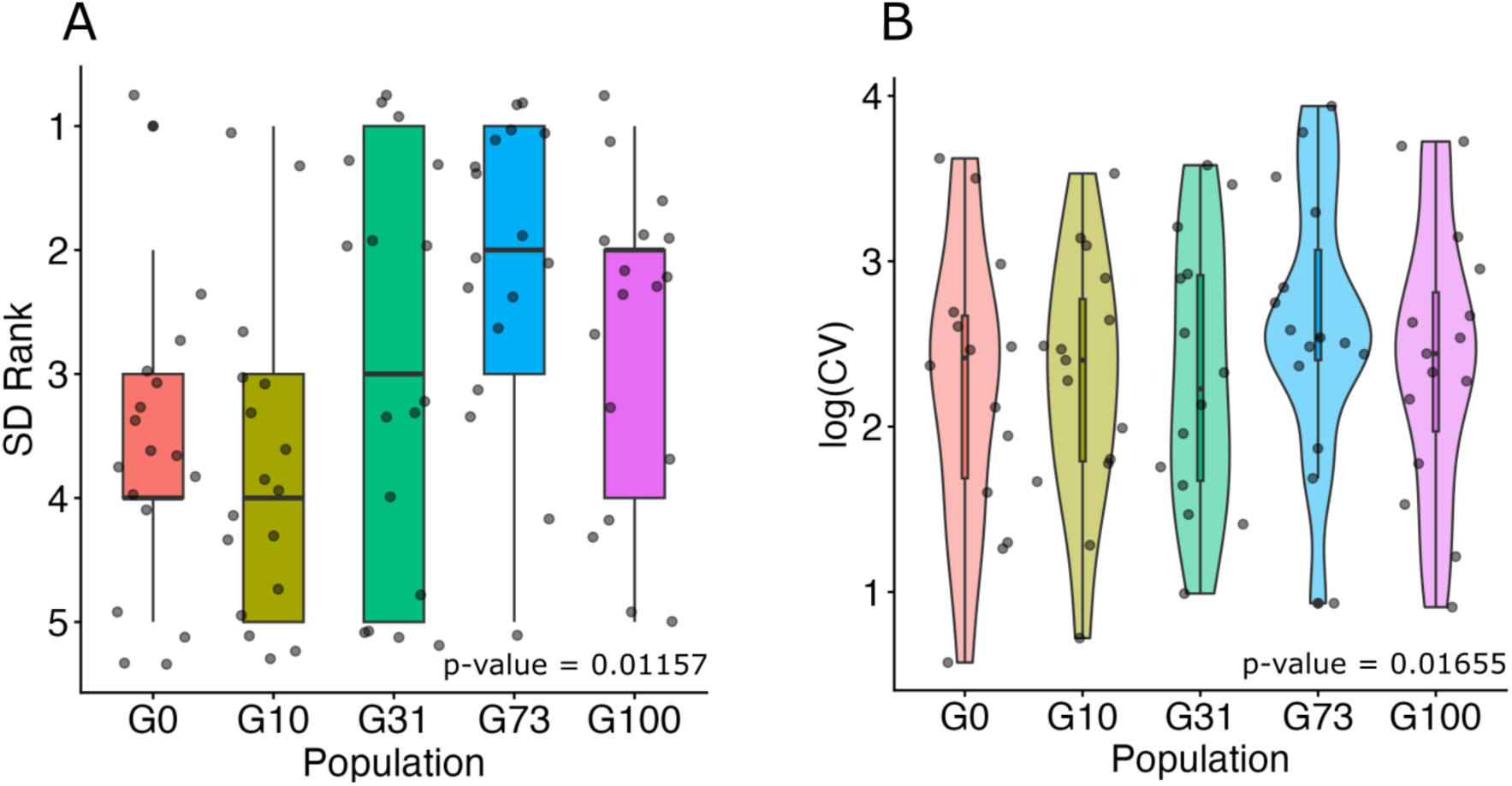
High TE content and insertional polymorphism increase phenotypic variation in females. A) Ranking of standard deviations (SD) across traits. Ranking differences were assessed using a Friedman test as indicated by the p-value in the plot. B) Logarithm of the coefficient of variation of the trait (CV) across the measured phenotypes. Each dot represents the CV for a single phenotype. Statistical comparisons between populations were performed using a linear mixed-effects model (LMER), the p-value of which is shown in the plot.

CV was again statistically higher for the G73 population compared to G10 (LMER p-value = 0.022), accompanied by a trend towards being higher than G0 (LMER p-value = 0.092) (Figure 5B). Both methods led to similar results, supporting the hypothesis that indeed G73 having a greater TE content associated with higher insertion polymorphism in the population generally expresses a wider range of variation of its phenotypes. G31 may not yet have accumulated enough TEs for this pattern to be observed. As for G100, although it carries a slightly higher number of insertions than G73, its phenotypic profile appears closer to that of G0 (Figure 5A). The particular profile of G100 may be explained by its TE abundance as assessed by DeviaTE (Weilguny and Kofler 2019). Indeed, we conducted an analysis of TE abundance per haplotype using a normalized coverage method with DeviaTE, which unlike TrEMOLO considers all TE-derived DNA sequences that align to the TE consensus provided, not only full-length insertions (Weilguny and Kofler 2019). This analysis revealed that G100 shares a similar overall TE abundance with G10 and G31, with values of 1263, 1257, and 1303, respectively. This result was consistent across three replicates of 35 X G100 subsampling. For reference, TE abundance in G73 was estimated at 1466, which is considerably higher than in the other four populations (Figure S5). Hence, this may explain why only G73 exhibits greater phenotypic variation. While such a hypothesis might partly explain the behaviour of G100, our understanding remains limited by the lack of transcriptomic data on TE and gene expression interactions, and on the overall epigenetic landscape.

This suggests that the amount of within-population TE polymorphism can yield significant phenotypic variability. This study provides evidence that TEs influence phenotypic variance and, for the first time, demonstrates this effect across an extensive diversity of phenotypes.

### Variation in TE content impacts the response to environmental changes

The previous sections have shed light on the contribution of overall TE composition to, respectively, inter- and intra-population phenotypic variation. Previous reports have described environment-dependent phenotypic variation mediated by specific TE insertions: for instance, a *roo* insertion acting as an immune-sensitive enhancer driving variation in survival to *P. entomophila* infection in *D. melanogaster* (Merenciano and González 2023). However, whether the global TE composition contributes to the response to environmental changes remains to be interrogated. To this end, four experiments testing environmental responses were performed on the five TE-divergent populations: longevity assays under two temperatures and four diets, oxidative stress resistance assays at two different oxidant concentrations, and a metabolic rate assay at three different temperatures. We adopted the reaction norms framework, including its associated statistical approaches, to characterise variation in phenotypes across environments. Although this framework traditionally relies on identical genotypes, working with populations issued from a parental isogenic line allows us to capture the environmental responses driven by insertional polymorphism. Briefly, the longevity under thermal stress varied non-linearly with TE load, dietary restriction had sex-specific effects and oxidative stress resistance (paraquat) showed population-specific slopes.

First, longevity was assessed in both males and females at 18°C and 22°C, as it is well established that poikilothermic organisms, such as *D. melanogaster*, experience significant modulation of lifespan in response to temperature changes, making lifespan a plastic trait (J. R. David and Capy 1988; Zwaan et al. 1992; Vermeulen and Bijlsma 2004). A two-step approach was used to analyse longevity. First, the estimated time when 80% of the flies died (LT80) was computed along with survival estimates under a probit distribution. Second, the data were analysed using a linear model (LMER) based on the time of death of each individual fly (see Material and Methods). Longevity decreased significantly with the increase in temperature in both sexes (Figure 6A) (females’ p-values: GLMER = 5.2e-6, LMER < 2.2e-16, males’ p-values: GLMER = 3.5e-12, LMER = 1.6e-14). In males and females, both LT80 values and LMER analysis support a strong effect of the interaction population by environment, suggesting that different populations and hence TE composition, experience different responses to temperature. The G31 population shows a rather drastic decrease in longevity with temperature increase (slope = -12.9), while G10’s does not show significant differences (slope = -1.5) (Table S3). These two cases illustrate the differences in the response to temperature changes. In males, reaction norms are overall less steep than in females (average slope = -3.5 in males vs. -7.8 in females), which indicates that males’ longevities are less impacted by temperature changes than females’ (Table S3); however, they still reveal variation in responses between the populations (Figure 6A). In both males and females, the environmental response appears to be more influenced by the TE composition rather than the overall TE content. Specifically, longevity does not follow a consistent pattern across generations of TE accumulation, suggesting that the composition of TEs, rather than their sheer number, plays a role in shaping the response to environmental changes.

**Figure 6:**
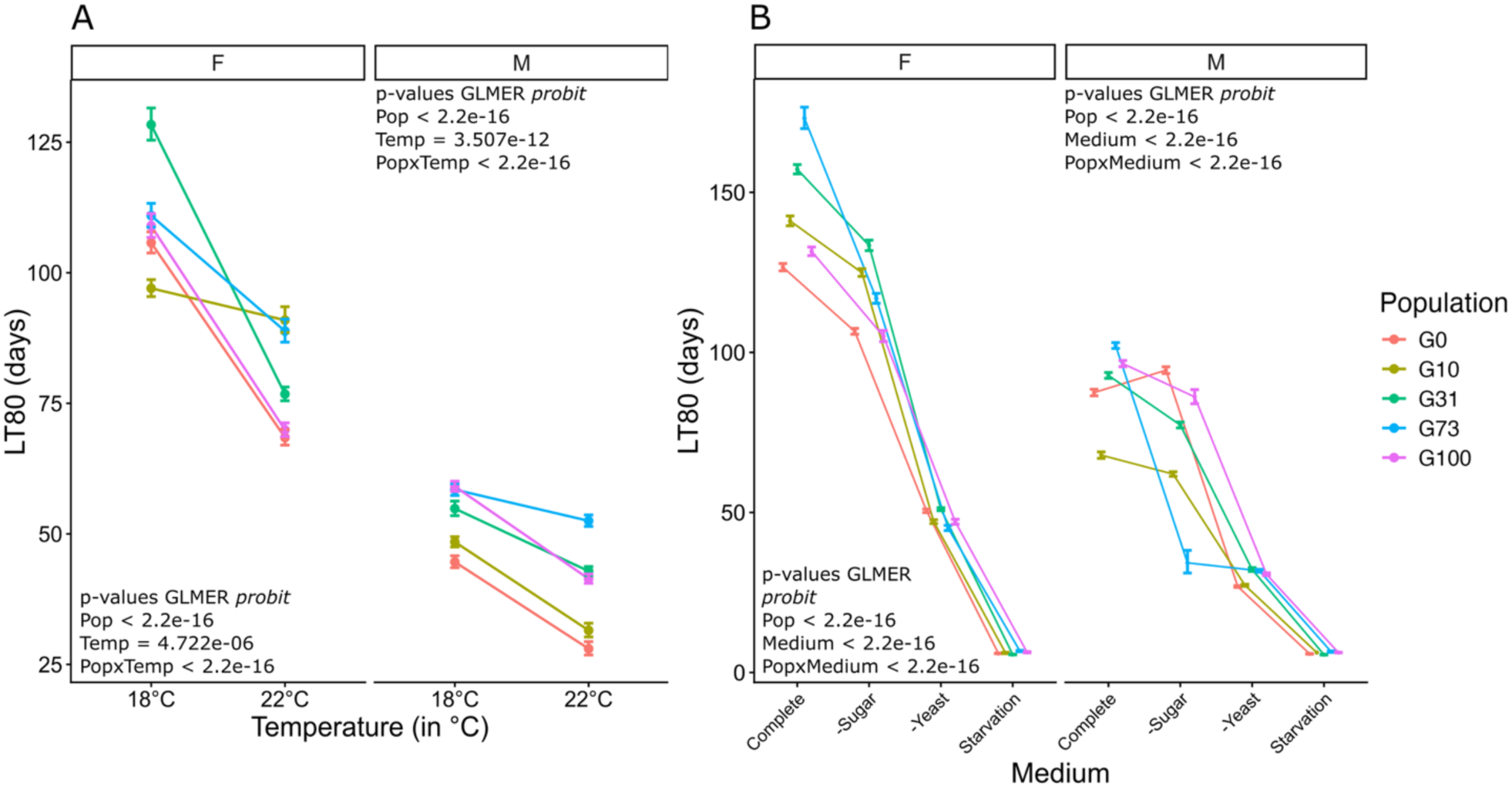
Longevity varies under variable temperatures and diets. A) Reaction norms of the LT80 for the females on the left and males on the right at variable temperature, namely 18°C and 22°C. B) Reaction norms of the LT80 values for females and males, fed on different media. The complete media represent a standard media in which the corn flour was removed, “-Sugar” is the complete media without the sugar, “-Yeast” is the complete media without the yeast and “Starvation” is a media with only water agar. F refers to the females and M to the males.

Measures of longevity at 18°C were also conducted on three different diets (Figure 6B). The first diet was composed of a complete medium containing a 5:4 ratio of proteins to carbohydrates, in the form of yeast and sugar. In the other diets, the medium was alternatively deprived of one or the other nutritional source, while in the last diet, the medium was deprived of all nutritional sources as a way of testing starvation resistance. In both males and females, we observed a strong population-by-environment effect (all p-values < 2.2e-16). LT80 values highlight the gradual decrease in lifespan as medium is deprived of nutritional value (Figure 6B). Interestingly, deprivation of sugar from the diet appears to have a greater impact on females than on most male populations, except for G73 (Figure 6B). In fact, where the decrease in lifespan is gradual with depletion in nutritional values in females, in males the deprivation in protein intake, by removal of yeast from the media, seems to have a greater effect than sugar deprivation (Figure 6B). The differences between males’ and females’ responses to dietary restrictions are an indicator of the sexual dimorphism and the variation in the nutritional needs. It was previously reported that female *D. melanogaster* lifespan is highly responsive to dietary sugar concentrations and more susceptible to their detrimental effects (Chandegra et al. 2017). Males are naturally shorter-lived, and their response to dietary restriction might be comparatively more subtle. Overall, strong interactions between populations and the environment are observed, suggesting the importance of TE composition in the response to changes in diet. However, the underlying reasons for the differences between the populations remain to be investigated.

Next, we investigated metabolic rate which provides a single point estimate for the energy expenditure needed to maintain fundamental organismal processes in a given environment (Lighton and Lighton 2021). Metabolic rate can be influenced by various biotic factors, such as body weight or age, as well as abiotic factors, including temperature, as tested here (Arking et al. 1996; Berrigan 1997; Van Voorhies 2009; Alruiz et al. 2023). As is expected in ectotherms, metabolic rates increased with temperature for both sexes (p-values = 4.72e-6 and < 2.2e-16 for females and males, respectively). Furthermore, metabolic rates differed between populations (GLMER p-values < 2.2e-16 for both sexes), but there was no significant interaction between the population and temperature, suggesting that all populations reacted similarly to the environmental change regardless of their TE composition and polymorphism (Figure 7A and 7B). Notably, the dry weight (included as a covariate in the model due to its positive correlation with metabolic rate (Kozłowski et al. 2020)) accounted for a significant portion of the variance in females but none in males. Conversely, the opposite trend was observed for the activity covariate, similarly included in the model for its relevant correlation with metabolic rate (Reinhold 1999) (Table S4 and S5). Videlier and collaborators observed a similar pattern when measuring the basal metabolic rate (BMR) in *D. melanogaster* (Videlier et al. 2019). They reported a positive correlation between activity covariate and BMR in males, whereas in females, BMR was either uncorrelated or even slightly negatively correlated with activity. Furthermore, a follow-up study on the genetic components highlighted sex-specific differences in the genetic architecture of locomotor activity and its impact on BMR (Videlier et al. 2021). This reflects the differences in energy investment between the sexes, as it is the case for gamete energy allocation, where females across all animal kingdoms invest more than males (Hayward and Gillooly 2011). Females are likely investing energy in reproduction, particularly in egg production, with larger females typically producing more eggs (Lefranc and Bundgaard 2000). In contrast, males’ reproductive investment relies more on their ability to find and court a mate (Asahina 2018). Therefore, males likely invest more energy in activity than females. These covariates reflect the sexual dimorphism in energy allocation. However, no interaction between TE composition and the environment is observed, as the reaction norms follow a comparable increase across the generations. This suggests that the effect of TEs may occur on very specific traits (as previously observed), rather than having an overall effect on the metabolism of the organism. It is worth noting that G0 tends to exhibit a higher metabolic rate in both males and females; given that metabolic rate summarizes many biological processes and that TE content may alter several of these processes, this observation may deserve further investigation.

**Figure 7:**
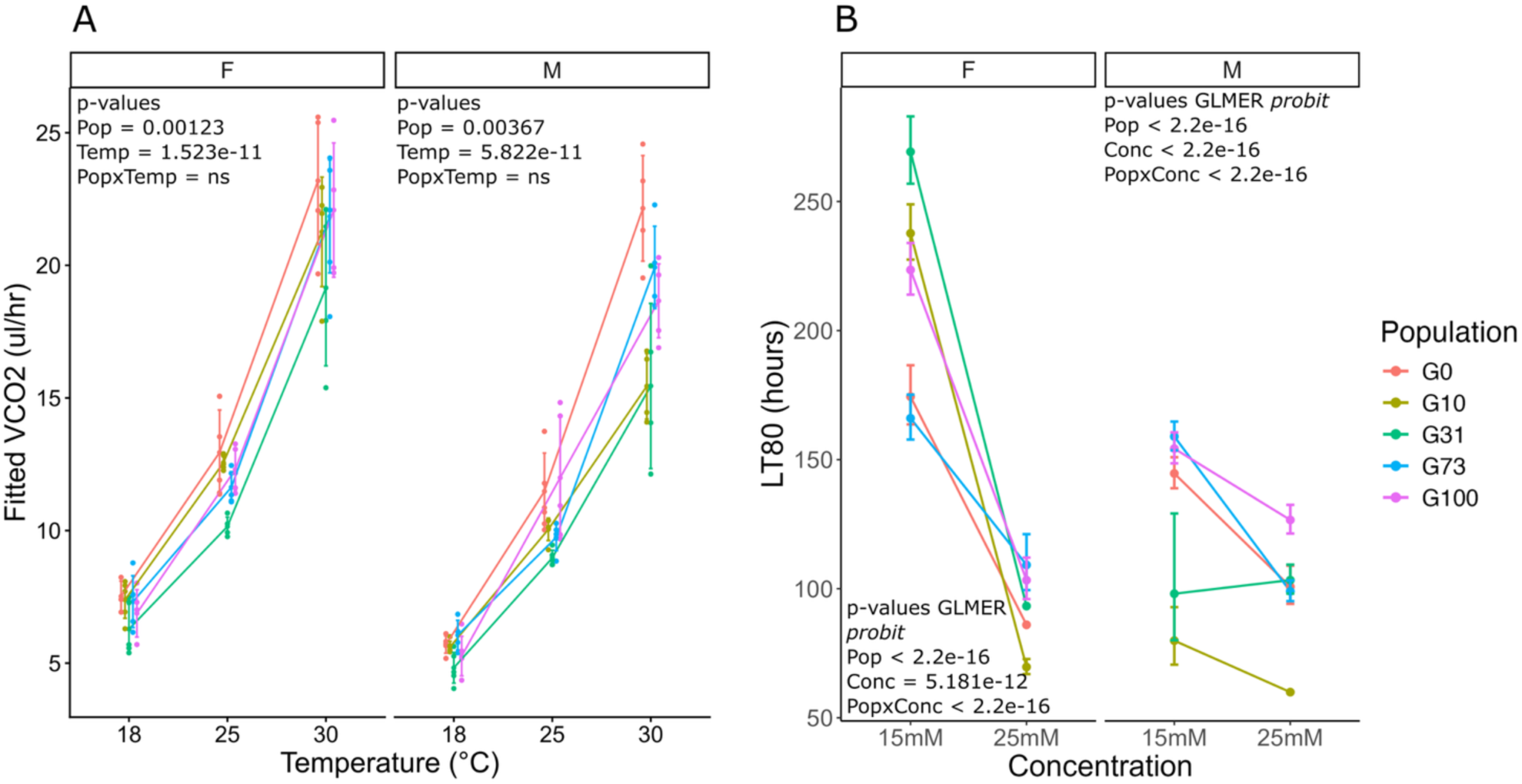
Metabolic rate and resistance to oxidative stress under variable conditions. A) Plot of the fitted values of the metabolic rate for the females (left panel) and males (right panel). Each condition is represented by five replicates of a pool of five flies. The fitted values account for the dry weight and the activity as covariants. B) Plot of the LT80 values under variable concentrations of the oxidative agent paraquat, for the females on the left and the males on the right. F refers to the females and M to the males.

Finally, the populations were exposed to two different paraquat concentrations, to measure the effect of different oxidative stress intensities. Exposure to the herbicide paraquat increases the formation of reactive oxygen species (ROS), known for their impact on longevity by causing cellular damage (Oleson et al. 2021; Shields et al. 2021; Costantini 2024). In both males and females, we report a significant interaction between TE composition and paraquat concentration (Figure 7B) (Table S6 and S7). Female survival is significantly reduced by the increase from 15 to 25 mM of paraquat (average slope = -12.2), whereas males’ hardly drops (average slope = -2.9) (Figure 7B, Table S8). Interestingly, females’ G73 survival difference from 15 to 25mM is significantly reduced compared to the other populations, with a slope of -5.7 (Figure 7B). This last observation seems to be tied to the fact that G73’s resistance to the lowest concentration of paraquat is significantly lower than that of the other populations, except for that of G0 (Figure 7B). Estimated survival probabilities from the GLMER model suggest that only females’ G0 and G100 chances of survival from one time point to the next do not change with paraquat concentrations. Additionally, at 25 mM, females G0 and G10 have significantly lower probability of survival than the other populations, echoing the results obtained for the longevities under different temperatures. G10 males have a strikingly low resistance to paraquat and especially at 25 mM concentration, with LT80 values around 60 h, for 100 h to 150 h in the other populations (Table S9).

Overall, all measured traits displayed variation in the environmental response, of which only metabolic rates measured under variable temperatures are not under the influence of an interaction between TE composition and environment (Table 3). These findings bring evidence for TE-driven environmental response in *D. melanogaster*. To determine the relative contribution of each variable (population, environment, and their interaction) to the observed phenotypes, we computed the Partial Eta^2^ for each variable (Figure 8). While the environmental effects were statistically significant, the associated effect sizes were sometimes subtle, as is the case for the longevity under variable temperatures, and the survival to different levels of oxidative stress. In the case of metabolic rate, however, the effect size showed a strong response to the environment, even though the statistical support was less pronounced. The discrepancies may lay in the fact that while the effect is statistically supported, the amplitude of the effect can be low (Sullivan and Feinn 2012; Dunkler et al. 2020). This can be explained by large sample sizes (as it is the case for the longevity measurements), whereas the reciprocal situation can be explained by small sample size.

**Figure 8:**
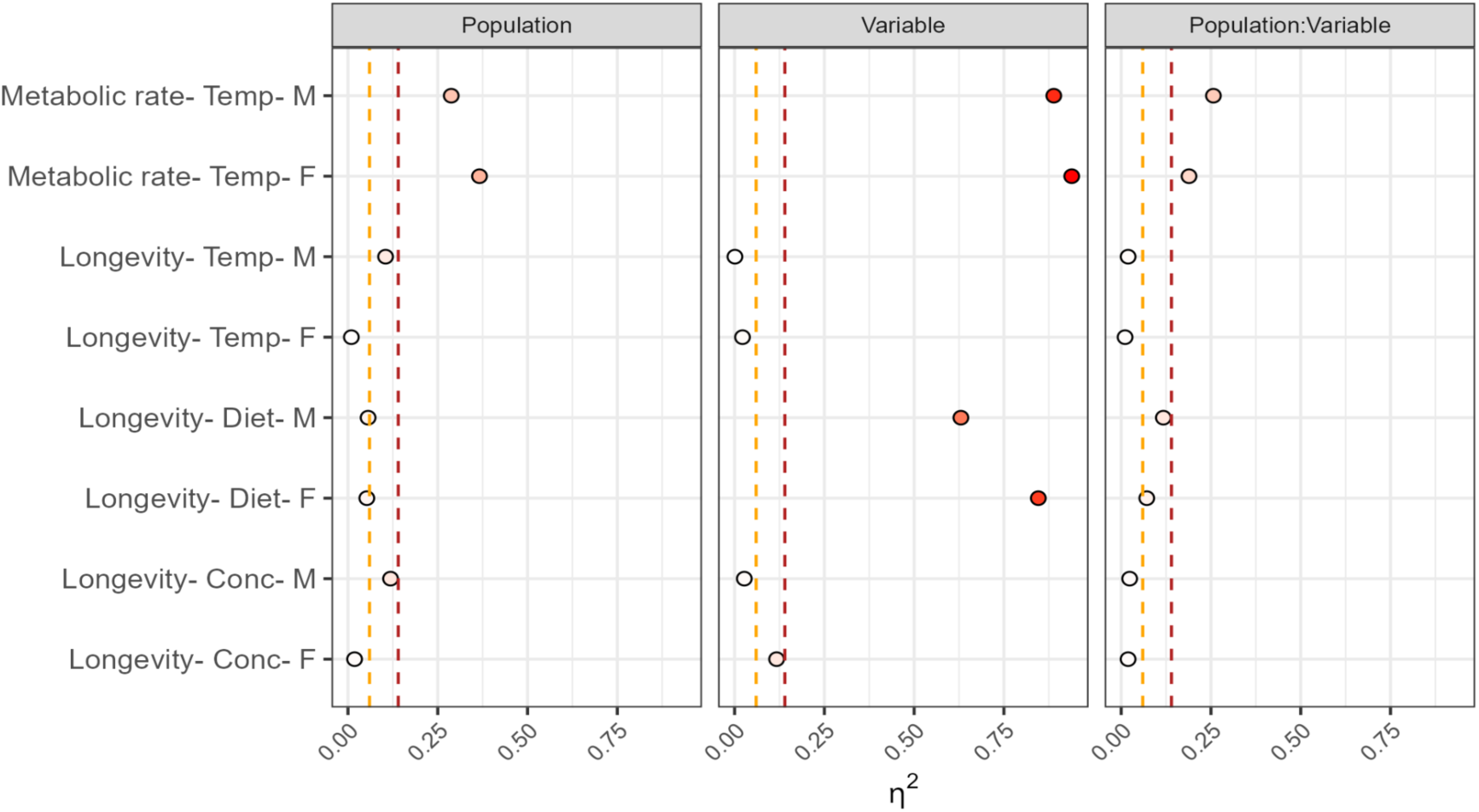
TE content contributes to differences in environmental responses of various traits across environmental conditions. This plot displays the effect size of the population, environmental variables (“Variable”) and their interaction on the phenotypes. The yellow vertical dashed line represents the threshold of medium effect and the red, the threshold of strong effect. “F”: females, “M”: males, “Temp”: temperature, “Conc” refers to the concentration of paraquat and “Diet” to the different media on which longevity was measured.

**Table 3:** Summary table of statistical analysis and results of the phenotypic measurement under variable environment. The “Phenotype” column indicates the trait assessed, “Measure” specifies how the trait was quantified, and “Environment” indicates the conditions under which the measurement was performed. “Sex” identifies whether the measurement was made on females (F) or males (M). “Population effect” denotes the statistical significance of differences among populations. “Interaction term” refers to the statistical significance of the effect of the interaction of the population with the environment. “Statistical test” specifies the model used to assess the population effect, including the linear mixed-effects models (LMER) and generalized linear mixed-effects models (GLMER). For the metabolic rate, the LMER model accounted for dry weight and activity as covariates. Statistical significance is depicted as follows: 0 ‘***’ 0.001 ‘**’ 0.01 ‘*’ 0.05 ‘.’ 0.1 ‘ns’ 1

| Phenotype | Measure | Environment | Sex | Population effect | Interaction term | Statistical test |
| --- | --- | --- | --- | --- | --- | --- |
| Longevity | Variable temperatures | Temperature | F | * | *** | LMER |
|  |  |  | M | *** | ** |  |
| Metabolic rate | CO2 emission | Temperature | F | ** | ns | LMER<br>(covariants:<br>dry weight<br>and activity) |
|  |  |  | M | ** | ns |  |
| Longevity |  | Diet | F | *** | *** | LMER |
|  |  |  | M | *** | *** |  |
| Oxidative stress |  | Paraquat concentration | F | ** | *** | LMER |
|  |  |  | M | *** | *** |  |

### Investigation into structural variants reveals candidate TE insertions

Acknowledging that global TE composition contributes both to phenotypic variation and to the response to environmental change, does not exclude the possibility of having phenotypic effects due to the impact of specific TE insertions. Hence, to detect candidate insertions involved in the occurrence of given phenotypes, two approaches were employed. First, we filtered TE insertions, across all families and classes, retaining those with a frequency ≥60% and unique to each population, thereby selecting insertions likely to have a population-level phenotypic impact. Second, we performed a genome-wide association analysis using Baypass (see Material and Methods).

From the first approach, a total of 10 putative candidate insertions, across the five populations, were identified through manual curation by considering insertions unique to each population with a frequency above or equal to 60% (see Material and Methods) (Table S10). Among them, three candidate insertions were *Doc* elements, consistent with its observed increase across generations. Three other candidate insertions belonged to the *roo* family, which has been reported to be especially active in these populations (Varoqui et al. 2025). Additionally, three candidate insertions belonged to the LTR class: a *Quasimodo* element, and one each from the two previously observed responsive families, *Zam* and *gtwin*. The insertions ensuing from TE families described to have markedly increase across the populations are of particular interest, as they are recent insertions and likely fully functional, thus retaining key regulatory or coding components.

From the second approach using Baypass in females (analysis in males yielded nearly identical results, since sex chromosomes were excluded, see Materials and Methods), comprising 843 insertions and the means and LT80 values of the 17 phenotypic measurements, five markers emerged with median Bayes Factor (BF) values above 14, thereby standing out as strong candidates (see Materials and Methods). Three of these makers had multiple significant associations to different traits. Of these candidates, one corresponded to a *roo* element, and one to the insertion of a *Doc* element. These are once again rather expected considering the dynamic nature of *roo*, as well as the notable increase in insertion of *Doc* in the present populations. The complementary manual curation of the Baypass output raised the list of candidates to 11 (see Material and Methods), with BF ranging from 8 to 20 (note that the same markers can have different BF values for each phenotype) (Table S11 and S12). Markers with high frequencies and with association to multiple phenotypes were part of the manually curated selection. Conversely, this curation also resulted in the removal of markers with strong association, but low frequencies and an absence of function described about the genes, with only two of the five markers with strong support kept in the final list of candidate insertions (Table 4). Likewise, non-repeat SVs for the females were submitted to the Baypass genome-scan. Out of the 290 cumulative non-repeat SVs, none emerged as a candidate (Table S13).

**Table 4:** Summary table of candidate insertions based on females’ data analysis through both manual and genome-their potential phenotypic effects. Following the chromosome and position of the insertion, the “Region” column indicates the genomic location of the TE in identity, “Population frequency” the insertion frequency across the TE-divergent populations in regard to G0, “Approac the identification, “G0 annotation” the reference gene annotation in the G0 genome, “Reported phenotypes (FlyBase)” th associated with the gene in FlyBase, and “Phenotype in the present study” the phenotypes observed in this study that ma *association detected by Baypass and Bayesian Factor above 14.

| Chromosome | start | end | TE | Region | Population frequency |  |  |  | Approach | GO annotation | Reported phenotypes (flybase) | Phenotype in the present study |
| --- | --- | --- | --- | --- | --- | --- | --- | --- | --- | --- | --- | --- |
|  |  |  |  |  | G10 | G31 | G73 | G100 |  |  |  |  |
| chr2R | 18047870 | 18047871 | con19_Quasimodo#LTR/Gypsy | Intron | 60 | 0 | 0 | 0 | Manual/Bypass | 5-HT1A | short-lived | LT80 (25mM paraquat), Dry weight* |
| chrX | 11594589 | 11594590 | con48_roo#LTR/Pao | Intron | 80 | 0 | 0 | 0 | Manual | Ptp10D |  |  |
| chr2L | 19750653 | 19750654 | con12_Doc#LINE/Jockey | Intron | 50 | 0 | 0 | 0 | Bypass | Lim3 | short-lived |  |
| chr3R | 875331 | 875332 | con12_Doc#LINE/Jockey | Intron | 0 | 100 | 0 | 0 | Manual/Bypass | Myo81F |  | Development time |
| chr3R | 939426 | 939427 | con12_Doc#LINE/Jockey | Intron | 0 | 90.9091 | 0 | 0 | Manual/Bypass | Myo81F |  | Development time |
| chr3R | 1614401 | 1614402 | con12_Doc#LINE/Jockey | Intron | 7.40741 | 0 | 21.2121 | 14.8148 | Bypass | Myo81F |  | Development time |
| chr2L | 2009581 | 2009582 | UnFUnCIUnAlig_RLX-incomp_FIN#LTR/Gypsy | 2kb upstream | 0 | 30.4348 | 0 | 0 | Bypass | CG42789 |  | LT80 (-Sugar&-Yeast), Starvation, Climbing, Dev, VCO2 (25&30C)* |
| chr4 | 886098 | 886099 | con48_roo#LTR/Pao | 5'UTR | 100 | 6.89655 | 0 | 0 | Bypass | Unc-13 | abnormal size | Dry weight |
| chrX | 219907 | 219908 | con12_Doc#LINE/Jockey | Intron | 0 | 80 | 0 | 0 | Manual | CG3777 |  |  |
| chrX | 4169729 | 4169730 | con48_roo#LTR/Pao | 3'UTR | 0 | 80 | 0 | 0 | Manual | CG15570 |  |  |
| chr4 | 1014820 | 1014826 | con16_blood#LTR/Gypsy | Intron | 0 | 90 | 0 | 2.5641 | Bypass | PlexA | abnormal starvation stress response | Starvation |
| chr3L | 12104738 | 12104739 | con12_Doc#LINE/Jockey | Downstream (<5kb) | 27.2727 | 0 | 52 | 0 | Bypass | Rbfox1 | abnormal locomotion, abnormal size | Locomotion, Dry weight, Oxidative stress |
| chr2L | 17674467 | 17674468 | con14_F-element#LINE/Jockey | 2kb upstream | 0 | 0 | 71.4286 | 0 | Manual | lncRNA:CR43802 | involved in spermatogenesis | Hatchability |
| chr3R | 21186346 | 21186347 | BDGP_ZAM#LTR/Gypsy | 2kb upstream | 0 | 0 | 66.6667 | 0 | Manual | CG34377 |  |  |
| chr3L | 20902911 | 20902912 | con2_gtwin#LTR/Gypsy | Intron | 0 | 0 | 0 | 69.2308 | Manual/Bypass | pip |  |  |
| chr3R | 27577719 | 27577720 | con48_roo#LTR/Pao | Intron | 0 | 0 | 0 | 64.2857 | Manual/Bypass | sima | abnormal stress response | Ctmax |

By integrating results from both approaches, we compiled a comprehensive set of 16 candidate TE insertions spread across the derived populations (G10, G31, G73 and G100) (Table 4). In particular, five insertions were identified by both methods and located near functionally relevant genes representing especially strong candidate loci, requiring investigation in future studies.

### Conclusions

Understanding the sources of phenotypic variation remains a central question in evolutionary biology, as does elucidating the impact of the effect of environmental changes on this variation. While the contribution of TEs to environmental responses is widely studied in plants (Negi et al. 2016; Joly-Lopez et al. 2017; Hirsch and Springer 2017; Ito 2022; Latzel et al. 2023; Pozzi et al. 2025), the field remains comparatively underexplored in metazoans. In insects in particular, TEs have been extensively investigated in the context of adaptive evolution (Casacuberta and González 2013; Gilbert et al. 2021; Cabral-de-Mello and Palacios-Gimenez 2024), yet their role in immediate environmental responses remains largely elusive and mostly theoretical (Capy et al. 2000; Piacentini et al. 2014; Horváth et al. 2017; Lanciano and Mirouze 2018; Marin et al. 2020; Pimpinelli and Piacentini 2020). This gap is not without reason: first, assessing the contribution of TEs to an organism’s response to environmental perturbation requires comparisons between individuals that are genetically identical except for their TE content, an exceedingly rare condition, if not unattainable, in natural populations. Secondly, tools for the analysis of TEs and their contribution to phenotypes are sparse, limiting the progression of this field. Few studies have previously highlighted the contribution of TEs to phenotypic variability. For instance, in *P-element*-burdened strains, the variance in traits related to fitness, viability, and fertility was observed to be higher than in the control strain (Mackay 1986). Alternatively, in a study comparing *Capsella rubella*, commonly called the pink shepherd’s-purse, and its sister species *C. grandiflora*, it was reported that although *C. rubella* exhibits reduced genetic variation due to a severe bottleneck event following speciation (Guo et al. 2009), it carries high levels of TE polymorphism compared to *C. grandiflora.* This turned out to be associated with a significant increase in transcriptomic variability near the *FLOWERING LOCUS C (FLC)* gene, a hotspot for *Helitron* insertions, which may explain the phenotypic divergence seen across natural populations (Niu et al. 2019). Although rare examples of TEs’ contribution to phenotypic variability are available, and especially in animals, there has been extensive theoretical work on the topic (Whitelaw and Martin 2001; Slotkin and Martienssen 2007; Rey et al. 2016; Chuong et al. 2017; Pimpinelli and Piacentini 2020; Catlin and Josephs 2022).

We took advantage of previously established *D. melanogaster* populations in which an increase in TE content was confirmed within a genetic background nearly devoid of other sources of genotypic variation, to evaluate the contribution of TEs to phenotypes and environmental responses (Barckmann et al. 2018). This study system provides an unprecedented opportunity to assess the role of TEs in shaping phenotypic variation. We first demonstrate that variation in TE composition – both in terms of content and insertional position – influences a broad range of phenotypic traits, including fertility-related traits, longevity, and behavioural traits such as climbing ability. More specifically, our results show that TE composition drives phenotypic divergence, increases intra-population variance, and modifies environment-dependent responses in Drosophila. As observed in the G73 population, while higher TE polymorphism increases phenotypic variability, it appears to come with a trade-off in fitness, as reflected in the impaired hatchability and viability of this population. This highlights the continuum of TEs’ influence, which can range from deleterious to neutral, and more rarely, to beneficial, depending on the environmental context. Such degraded fitness may have been detrimental to the point of selecting for an alternative TE control mechanism during the induction of TE derepression, resulting in a decrease in TE accumulation as observed in G100. While the underlying mechanisms remain unresolved, the TE content of the G100 seems to have reached a plateau, as evidenced by its phenotypic profile highly resembling that of G0’s. While these findings support the notion that an excessive TE load can be detrimental, it has also been shown that a complete absence of TEs may be disadvantageous. Cranz-Mileva and colleagues (2024) demonstrated that in *Schizosaccharomyces pombe*, strains completely depleted of the LTR retrotransposon Tf2 exhibited reduced fitness, whereas strains retaining TEs conferred a positive contribution to the host (Cranz-Mileva et al. 2024).

The assessment of the populations’ responses to environmental changes revealed that three of the four traits tested, namely longevity under different temperatures, under different diets and oxidative stress resistance, point to a divergence in environmental responses between the TE-divergent populations. The strength of this study lies in the diversity of assays, which provides robust support for the role of TEs in environmentally-induced phenotypic variation. To our knowledge, this is the first study to investigate the contribution of TEs to the response of environmental changes, across such a broad range of phenotypes and environmental conditions in insects, offering empirical evidence that bridges longstanding theoretical predictions with experimental validation. This further supports the role of TEs in shaping not only phenotypic variation, but also in contributing to environmentally-induced variation. In this context, TEs are standing genetic variants whose phenotypic effects may remain unexpressed under standard conditions but can be revealed following environmental perturbations, thereby being prominent representatives of cryptic genetic variants (CGVs) (Gibson and Dworkin 2004; Ledón-Rettig et al. 2014; Paaby and Rockman 2014; Schneider and Meyer 2017; Pfennig 2021). Because their effects are environmentally inducible, CGVs constitute intriguing drivers of phenotypic diversity, whereby identical individuals may exhibit distinct CGV effects across environments. Hence, CGVs are influential contributors to evolutionary potential, of which TEs are often an overlooked component. Indeed, the demonstration of TE-driven inter- and intra-population phenotypic variation, and more importantly of environmentally induced phenotypic variation, supports the contribution of TEs to generating phenotypic diversity and promoting the emergence of selectable TE-driven traits. This observation appears to be in conflict with the traditional view that TEs accumulate primarily in genomic positions where their impact is neutral, while deleterious insertions are rapidly removed by selection. Consequently, our results interrogate the extent to which TE insertions are truly phenotypically neutral, and how this relationship shifts as total TE content increases. To our knowledge, such empirical evidence has never been reported under comparably strict experimental conditions in any metazoan system. Moving beyond population-level studies, the establishment of isogenic lines carrying variable TE compositions but representing identical TE loads for each of the TE-divergent populations would allow us to test not only the effect of TE load, but also the phenotypic consequences of varying TE cohorts, opening avenues for investigating the mechanistic basis of phenotypic plasticity.

## Material and methods

### 1. Fly husbandry

A *D. melanogaster* line was previously modified to suppress the piRNA pathway in charge of TE regulation (Barckmann et al. 2018). Since a constitutive piRNA knockdown is lethal, a system was engineered to target ovarian somatic cells with a heat-induced piRNA KD. Briefly, the fly line carries three components: (i) a GAL4 UAS-activator driven by the follicle cell-specific *traffic jam* (tj)-promoter (tj-GAL4), (ii) a UAS-short-hairpin(sh)-piwi that induces *Piwi* RNAi and (iii) an ubiquitously expressed thermo-sensitive GAL4-inhibitor, GAL80ts (Barckmann et al. 2018). The initial isogenic population (G0) was repeatedly exposed to temperature changes while maintaining a population size of around 500 progenitors in standard medium-filled bottles, to reduce genetic drift. The consequence is the occurrence of independent transposition events in each individual of the offspring, leading to insertion polymorphism in a constant genetic background (Barckmann et al. 2018). In total, five populations harbouring different TE content and insertions were generated (G0, G10, G31, G73, G100). G0 is the parent population, and all subsequent shifts are named GX, where X is the number of generations that have been shifted. Hence, G10 corresponds to 10 generations of shifts (10 generations of piRNA KD) and therefore of TE accumulation. All gifted populations were naturally infected by the endosymbiont *Wolbachia*. Flies were maintained at 18°C in 12-hour light cycles, with 65% relative humidity, in vials with standard medium (0.2 g agar, 1.46 g maize flour, 1.53 g yeast, 0.08 g Nipagine, 0.63 mL ethanol, 12 mL water), with a density of approximately 100 progenitors at each generation. Each population was maintained in five vials per generation, which were frequently shuffled to avoid vial effects. All experimental procedures used flies reared and eggs collected under the environmental conditions described above. Adult flies were only subjected to environmental variations after completing development, as detailed in the respective methods sections.

### 2. DNA extraction and sequencing

DNA extractions were performed on 200 embryos per population using a homemade protocol designed to yield long DNA fragments. Briefly, tissues are homogenized in STE buffer (sodium-Tris-EDTA), to which is added 10% SDS (sodium dodecyl sulfate). The mix was incubated for 2 h 30 min with proteinase K at 55°C. The samples were then treated with RNase for 1 h at 37°C, and the DNA was precipitated with phenol/chloroform/isoamyl alcohol and purified by isopropanol and ethanol washing steps. The DNA was eluted in 35 µL of TE elution buffer (10 mM Tris-HCl (pH 8.0) and 0.1 mM EDTA) and stored at 4°C until library construction and long read sequencing. These steps were performed at the PSI sequencing platform at the Institut de Génomique Fonctionnelle de Lyon (IGFL). The DNA extracts were quantified using Qubit (dsDNA 1X HS Assay Kit, Thermofisher), revealing an average DNA concentration of 40 ng/µl. The distribution of DNA fragment lengths was verified by TapeStation 4150 (Agilent) on a Genomic ScreenTape, confirming high molecular weight DNA for all samples, with a maximum peak observed around 30 kb. The Oxford Nanopore Technologies (ONT) libraries were constructed using the ONT Native Barcoding Kit 24 V14 (SQK-NBD114.24), starting with 400 ng of DNA for each sample and following the ONT protocol recommendations. The pooled barcoded libraries were sequenced for 96 hours on a single run with a R10.4.1 flow cell (FLO-PROM114M) and PromethION 2 Solo sequencer with MinKNOW software version 23.11.7. To improve read quality, raw reads (pod5) were subjected to re-basecalling post-run using the Super Accurate basecaller (dna_r10.4.1_e8.2_400bps_5khz_sup.cfg; Dorado v7.2.13). Around 13M reads were generated in total, among which more than 11M had a quality above Q10, representing 53 Gbases sequenced of the best quality.

### 3. Genome assembly and gene and TE annotations

Raw reads were filtered for quality above Q10, and subsequently used to infer coverage and general statistics using the following tools: Nanoplot (v 1.42.0) (De Coster and Rademakers 2023), Cramino (v 0.14.5) (De Coster and Rademakers 2023), and Qualimap (v 2.3) (Okonechnikov et al. 2016). G0 was assembled from average coverage nearing 50 X. The genomic assembly was performed with Flye (v 2.9.3-b1797) using standard parameters except for the --nano-raw option (Kolmogorov et al. 2019), followed by four rounds of Racon (v 1.5.0) with minimap2 (v 2.29-r1283) conducted using the parameters for ONT data (Vaser et al. 2017; Li 2021). A step of purging was performed with Purge Haplotigs (v1.1.2) and Samtools (v 1.6) (Roach et al. 2018; Danecek et al. 2021). Finally, scaffolding was performed using RagTag (v 2.1.0) (Alonge et al. 2022) with default parameters using the dm6 version of *D. melanogaster* as the reference genome (https://hgdownload.soe.ucsc.edu/goldenPath/dm6/bigZips/). The Benchmarking Universal Single-Copy Orthologs (BUSCO, v 5.7.1) analysis was performed on the assembly using the Arthropoda reference gene set (Manni et al. 2021) (Table S14). The Quast pipeline (v 5.3.0) was also ran providing assembly parameters to evaluate assembly quality (Mikheenko et al. 2018) (Table S14). The G0 assembly was annotated using Liftoff (v 1.6.3) with Dm6 annotation (https://hgdownload.soe.ucsc.edu/goldenPath/dm6/bigZips/genes/dm6.ncbiRefSeq.gtf.gz) as reference (Shumate and Salzberg 2021). RepeatMasker (v 4.1.5) was run on the G0 assembly using the *D. melanogaster* Manually Curated TE library (MCTE) previously published (Smit et al. 2021; Rech et al. 2022).

### 4. Transposable element analysis

Rarefaction curves were generated to assess TE detection power using a logarithmic prediction model using the built-in Stat package R 4.4.0 (Bolar 2019; R Core Team 2021). TrEMOLO (Mohamed et al. 2023) was first run on subsets of reads, with coverage ranging from 5 X to the maximum coverage available for each population with 10 X increments. The number of TE insertions detected at each coverage level was used to model the expected number of insertions using a logarithmic function, allowing prediction of TE discovery saturation as coverage approaches the sequencing of all haplotypes. Downstream analyses were computed on 35 X coverage subsamples of the raw reads for all populations generated using Rasusa (v 2.0.0) (Hall 2022). TE variants and frequencies were obtained with TrEMOLO using default parameters of the OUTSIDER module and G0 as reference genome and the *D. melanogaster* MCTE TE library previously published (Rech et al. 2022; Mohamed et al. 2023). Using the logarithmic extrapolation of the TrEMOLO output, we estimate that at 35 X coverage, approximately 50% of TE variants are detected in each of the five populations (G10=49.74%, G31=53.238%, G73=52.38%, and G100 = 51.60%) (Figure S6). TE insertional variants were filtered from the TrEMOLO output file “TE_INFOS.bed”, and no_name contigs were removed for downstream analysis. Differences in average TE insertion frequency among populations were tested using the Kruskal-Wallis test in R (v 4.4.0) (R Core Team 2021). A shared insertion matrix was generated using a custom script, based on 100 bp windows upstream and downstream of the insertion site detected by TrEMOLO. Cases of ambiguous sharing where different TE class or family were detected at the same insertion site across the populations, were manually curated: insertions were merged if the same class/family/TE was present in the majority of the population or marked as unresolved if no consistent pattern could be identified. Additionally, when multiple insertions within the same population were found at the same position, the same criteria were applied, and their frequencies were summed. Plotting of the insertions per class and families was performed in R 4.4.0 (R Core Team 2021). DeviaTE (v2.2.1) was run on the subsampled data using default settings and three normalization single-copy genes: *Act5*, *Piwi*, and *Rpl32* (Weilguny and Kofler, 2019). The software aligns long reads to the TE library and estimates the TE insertion number for each TE family per haploid genome. The sum of high-quality (>Q15) coverage estimates was used as a proxy for TE load in each population.

### 5. Non-repeat structural variant analysis

The genome assemblies of all subsequent populations (G10, G31, G73, and G100) were generated following the same protocol used for the G0 assembly (see Materials and Methods: Genome assembly). The sequencing coverage of G10 was approximately 72 X, while G31, G73, and G100 had coverages of 43 X, 47 X, and 52 X, respectively. Structural variants (SVs) ≥100 bp (insertions and deletions) were detected using the GraffiTE pipeline with the *--genotype* parameter, comparing each assembly to the G0 reference genome (Groza et al. 2024). GraffiTE SVs were masked using RepeatMasker (v 4.1.5) to obtain non-repeat SV, specifically ruling out TEs and microsatellites (Smit et al. 2021). The GraffiTE pipeline was then diverted to align reads to the SVs and estimate their frequencies. The SVs found on “no_name” contigs were removed from subsequent analysis. SV length distributions and frequencies were plotted in R (v 4.4.0) (R Core Team 2021; Groza et al. 2024). SVs were intersected to determine which variants were shared among populations. An SV matrix was generated using a custom script. Cases of divergence in SV sequences at nearby positions were manually curated: SV sequences that were identical over at least 2/3 were merged into a single SV, while all others were kept separate. Frequency and size distributions were tested for patterns of divergence between populations using a Kruskal-Wallis statistical test.

### 6. Viability and development time

To measure viability of the egg-to-adult survival (Gasser et al. 2000), 150 three-day-old female-male couples (sexed and counted under CO2 anaesthesia) per population were placed in a small Dutscher egg-laying chamber (product reference: 789066). Flies were allowed to lay eggs for half a day. From these egg laying chambers; five replicates of 40 eggs were collected on small regular fly-food caps positioned in vials containing the standard media (Figure S7A). Eggs were collected under a binocular microscope, using the tip of a scalpel to scoop the eggs out of the media. The resulting vials were place in the climatic chamber maintaining the rearing conditions as previously described in Methods:Fly husbandry. Hatched flies were counted daily, from the first emergence to the last. Viability was computed as the proportion of adults emerging from the initial 40 collected eggs. Data were analysed under R software (v 4.4.0) using generalized linear mixed-effects model (GLMM) with binomial distribution using the Lme4 package (v 1.1-36) (R Core Team 2021; Bates et al. 2025). ANOVA Chi square test was applied to assess the fixed effect. McFadden’s R^2^ was estimated with the package Jtools (v 2.3.0) (Long 2024). From these data, the development time was also assessed, for each egg counted for its viability was also recorded the time until emergence of the adults. Using this data, a cumulative percentage of flies emerged each day since oviposition was calculated from the total number of eggs tested per replicate (five replicates of 40 eggs per condition). The comparative statistical analysis of the emergence curves between populations was performed using a Log Rank test in R (v 4.4.0) (R Core Team 2021). Development time was also analysed using ANOVA on the time at which each fly has emerged in R (v 4.4.0, Stat built-in package) (Bolar 2019; R Core Team 2021). Partial Eta^2^ was calculated using package Effectsize (v 1.0.1) (Ben Shachar et al. 2020).

### 7. Embryonic hatchability

To measure hatchability (the proportion of eggs from which a larva emerges (J. David et al. 1975)), 150 three-day-old female-male couples per population were placed in a small Dutscher egg-laying chamber (product reference: 789066). Flies were allowed to lay eggs for half a day under the previously described standard rearing conditions. Eggs were collected using a fine brush and water, and filtered through a fine mesh. Eggs remained dipped in water while five replicates of 50 eggs were collected and aligned on a black piece of Canson 160 g/m² paper (Figure S7B). The vials were placed in the climatic chamber maintaining the rearing conditions as previously described. Two days after sampling, the number of emerged eggs were counted under binoculars. Data were analysed under R software (v 4.4.0) using generalized linear mixed-effects model (GLMM) with binomial distribution using the Lme4 package (v 1.1-36) (R Core Team 2021; Bates et al. 2025). ANOVA Chi square test was applied to assess the fixed effect. McFadden’s R^2^ was estimated with the package Jtools (v 2.3.0) (Long 2024).

### 8. Dry weight

To conduct dry weight measurements, density-controlled vials were prepared. Specifically, 150 3-day-old female-male pairs per population were placed in a small Dutscher egg-laying chamber (product reference: 789066), with flies sexed and counted under CO₂ anaesthesia. Flies were allowed to lay eggs for half a day under standard rearing conditions. From each chamber, five replicates of 40 eggs were collected and transferred into vials containing standard fly medium. The vials were maintained in the climatic chamber under the rearing conditions previously described. Emerging adults were aged for five days, then sexed and pooled into groups of five. The flies were frozen at -20 °C overnight and subsequently dried in an oven at 75 °C for 24 hours. The dry weight data were analysed using ANOVA with R (v 4.4.0, Stat built-in package) (Bolar 2019; R Core Team 2021). Partial Eta^2^ was calculated using package Effectsize (v 1.0.1) (Ben Shachar et al. 2020).

### 9. Climbing assays

A pool of ten 3-to-5-day old flies was placed in pre-washed vials (2 cm in diameter, 9.5 cm in length), which had been dried and left to air out to eliminate any residual plastic odour. The vial was divided into three predefined zones: the first zone extended from the bottom to a height of 2 cm, the second from 2 cm above the bottom to 2 cm below the top, and the third from 2 cm below the top to the top of the vial. Prior to the trial, flies were placed in the tubes using a mouth aspirator and allowed to acclimate for 30 minutes. The flies were gently tapped three times to ensure they all fell to the bottom. After a two-minute trial, the number of flies in each zone was recorded. This procedure was repeated in batches of three experimental sessions, with each population and sex represented in every batch, resulting in six biological replicates (totalling 60 flies per condition). Taking the count of flies reaching the top of the tube as success, and the ones in the rest of the tubes as failure (middle and bottom zones), the statistical analysis was conducted in R (v 4.4.0) using a generalized linear mixed-effects model (GLMM) with a binomial distribution (R Core Team 2021; Bates et al. 2025). ANOVA was applied to assess fixed effects, using a Chi-square test. McFadden’s R^2^ was estimated with the package Jtools (v 2.3.0) (Long 2024).

### 10. Thermal tolerance

Critical thermal maximum (CTmax) was assessed for 5-to-7-day old females under controlled temperature ramping in a water bath, based on the same system as Riddell and collaborators (Riddell et al. 2023). Insects were placed in small vials inside a glass chamber. Water was heated with an aquarium water heater (Juwel Aquarium Aqua Heat 300) and two aquarium water pumps (Aqueon Circulation Pump 500) attached to it that continuously circulated warming water with an increasing rate of 0.6°C per minute, starting from 20°C until all flies entered a coma, typically around 40°C. The temperature was continuously monitored in each of the six chambers using type-T thermocouples (PT-6, ThermoWorks) embedded in the chambers and connected to an eight-channel thermocouple data logger (TC-08, Pico Technology) to ensure consistency across trials. Data were collected from six pools of five female flies each, totalling 30 individuals per condition. Statistical analysis was performed in R (v 4.4.0) using a linear mixed-effects model to account for variation between trials and arenas (R Core Team 2021; Christensen [2018] 2025). Post-hoc pairwise comparisons of CTmax across populations were conducted using estimated marginal means (EMMs), with Tukey adjustment for multiple comparisons. Partial Eta2 was calculated using package Effectsize (v 1.0.1) (Ben Shachar et al. 2020).

### 11. Longevity

Ten replicates of pools of 9 to 12 virgin females or males were placed in vials containing standard fly food, for each of the five populations (G0, G10, G31, G73, G100) (Table S15). Virgin flies were collected in the afternoon, after the vials were emptied in the morning, and were sexed under CO2 anaesthesia. All longevity replicates for all populations were established within one week, therefore using flies from a single generation. In addition, flies were placed on the same batch of standard medium. During the experiment, the vials were changed once every week to prevent low-quality media from impacting the experimental outcome. The cumulative number of dead flies was counted every 24 hours on weekdays. Survival curves and Log Rank test pairwise comparisons were generated with the Survminer (v 0.5.0) and Survival (v 3.5-8) packages in R (v 4.4.0) (Kassambara et al. 2021; Therneau et al. 2024). This experiment was conducted under 65% humidity, both at 18° C and 22°C. The time when 80% of the population died (LT80) was estimated using the Ecotox (v 1.4.4) package in R (v 4.4.0) (Wheeler et al. 2006; R Core Team 2021). The slopes were calculated from a linear model of LT80 *vs.* temperature. Analysis of the survival chances was performed using a generalized linear model (GLMER) and probit distribution with replicates as random effects. Type III ANOVA was applied to assess the fixed effects, and EMMs were adjusted using the Bonferroni adjustment method to account for multiple comparisons. Additionally, a linear comparative analysis of longevity at 18°C *vs.* 22°C was performed using a linear mixed-effects model with the lmerTest (v 3.1-3) package (Christensen [2018] 2025). Type III ANOVA was also applied to assess the fixed effects, and EMMs were adjusted using the Bonferroni adjustment method to account for multiple comparisons. Partial Eta^2^ were calculated using package Effectsize (v 1.0.1) (Ben Shachar et al. 2020).

### 12. Longevity with dietary restriction

Ten replicates of pools of 9 to 12 virgin females or males were prepared for each of the diet-specific longevity tests, except for the complete medium and the sugar deprived conditions, for which five replicates were conducted. Replicates were conducted in two separate batches of five, ensuring that each population was represented in both experimental batches (Table S16). Virgin females and males, collected as previously described, were transferred to the experimental media after ageing for four days on standard medium. All experimental media contained nipagin (Methyl 4-hydroxybenzoate) as a preservative. The diets were derived from the standard rearing medium, maintaining a sugar-to-yeast ratio of 5:4 in the complete medium, with corn flour removed. The yeast-deprived medium was based on the complete medium formulation with yeast omitted, and the sugar-depleted medium was similarly prepared by omitting sugar. The starvation medium consisted of agar and nipagin only as a preservative. All vials were maintained in a climatic chamber where temperature and humidity were identical to the breading conditions. Mortality was recorded three times per day (at 9 am, 1 pm, and 5 pm) every day until all flies had died. Time of death was then converted into days for analysis. Statistical analysis was performed using the same workflow as for longevity assays, only temperature was replaced by medium. Additionally, the starvation data were analysed separately as an independent trait, using a similar statistical approach.

### 13. Survival to oxidative stress

Resistance to oxidative stress was assessed using 15 mM and 25 mM concentrations of paraquat. Ten replicates of pools of 9 to 12 virgin females or males were first aged for four days on standard rearing medium, under controlled temperature and humidity as previously described, and then transferred to media containing either concentration of paraquat (Table S17). Additionally, five replicate pools of approximately ten virgin females or males were maintained on standard medium as controls. The number of dead flies was recorded three times per day (at 9 am, 1 pm, and 5 pm) every day until all flies died. Time of death was then converted into days for analysis. Statistical analysis was performed using the same workflow as the longevity assays, only temperature was replaced by concentration. The longevity of the mock condition was analysed and no significant differences between the populations were reported.

### 14. Metabolic rate

Metabolic rate measurements were obtained with a MAVEn™ flow-through respirometry device from Sable Systems. During trials, sixteen 5mL chambers containing 5 flies from each combination of sex and population were measured for CO2 production (VCO2) with a LiCor 850 gas analyzer as a proxy for metabolic rate, which is common in small insects (Lighton and Lighton 2021). Five replicates (chambers) per condition were conducted, with 3 (repeated) measures taken for each chamber. Chambers were measured at one of three experimental temperatures: 18°C, 25°C and 30°C. All trials occurred in the dark to normalize light exposure and reduce activity; however, system specific activity measurements for each chamber were additionally recorded using infrared-light detectors. After trials, flies from each chamber were frozen at -20°C and dried for 24 h at 75°C for subsequent weighing. VCO2 log-transformed data were analysed with linear mixed effect models in R (v 4.4.0) with temperature and population and their interaction as fixed effects, dry weight and activity as covariates, and chamber identification and trial as random effects (R Core Team 2021; Christensen [2018] 2025). Type II ANOVA was also applied to assess the fixed effects, and EMMs were adjusted using the Bonferroni adjustment method to account for multiple comparisons. Partial Eta^2^ were calculated using package Effectsize (v 1.0.1) (Ben Shachar et al. 2020).

### 15. Evaluation of the phenotypic variation

To assess differences in phenotypic trait variability across populations, two complementary methods were employed. First, the raw standard deviation was calculated for each trait within each population and used as a proxy to phenotypic variation. These values were analysed using Friedman’s rank test to evaluate whether variation in TE content explains phenotypic variability. As a complementary approach, the coefficient of variation (CV) was also computed for each trait (Colwell 1974; Burgess and Marshall 2014; Morgante et al. 2015). CV values were log-transformed (ln(CV)), and a linear mixed-effects model (LMER) was applied using the lmerTest package (v3.1-3), with the trait included as a random effect (R Core Team 2021; Christensen [2018] 2025). EMMs were adjusted for multiple comparisons using the Bonferroni correction, enabling population-level pairwise comparisons.

### 16. Linking TE and non-repeat structural variants with phenotypes

The first approach consisted in investigating insertions present in high frequency in only one or a few populations experiencing specific phenotypes, so called *a priori*-approach. This method was reinforced by the investigation of insertions positioned in candidate regions, such as insertions found in 3’ and 5’ UTRs, exons and promoters. Using the insertion frequencies from TrEMOLO OUTSIDER module output (Mohamed et al. 2023), TE insertions were filtered based on frequency, using a threshold of ≥60% prevalence to identify variants likely to have a phenotypic impact at the population level. Insertions were then further filtered to retain only those present in a single population (within 100 bp windows). These were then intersected with the different gene regions of G0. TE insertion positions were cross-referenced with Flybase (https://flybase.org/) and manually curated to retain those with reported phenotypes relevant to the traits under investigation in this study. For the second approach, a genome-wide association analysis was conducted using the Baypass tool, employed following the Bayes Factor (BF) (expressed in deciban (dB)) methods from Mérel and collaborators (Mérel et al. 2021). Indeed, Baypass (v 2.41) was used to identify TE insertions involved in insertion-phenotype associations (Gautier 2015). By leveraging insertion frequencies obtained from TrEMOLO, we inferred the allelic frequencies of each insertion for each population and analysed the data using the Baypass pipeline. We then identified loci (*i.e.* insertions) with high BF values, indicating strong associations with phenotypic variation within a Bayesian framework. Since sequencing was conducted on pooled embryos with an unknown female-to-male ratio, we excluded insertions on sex chromosomes and only analysed autosomal TE insertions. Five iterations of Baypass were conducted, and the median of the five BF was taken as the indicator of strong associations with phenotypic variation within a Bayesian framework. Insertions with BF values greater than 14 were considered strong candidate insertions in accordance with Jeffrey’s rule (Jeffreys 1948). In addition, an *a priori* manual curation was conducted to refine candidate selection for insertions with BF values below this threshold, considering their frequencies, genomic positions, and associated phenotypes as reported in FlyBase. Since Baypass utilizes phenotypic data, male and female datasets were analysed separately. The genome-scan approach using Baypass was applied to the non-repeat SVs following the same approach.

### Supplementary data

All supplementary data are compiled in a Supplementary Material document and include supplementary figures (Figures S1 to S7) and tables (Tables S1 to S17). Large tables are provided as separate supplementary files (Tables S11, S12, and S13), whose legends are described in the document.

### Data availability

The raw phenotypic data and the processed output of the transposable element analysis (downstream of the cluster-implemented pipelines TrEMOLO, GraffiTE, DeviaTE, and Baypass) and the corresponding R Markdown scripts are available at https://doi.org/10.5281/zenodo.22070083.

## Acknowledgments

We wish to acknowledge the insightful discussions about TE analysis with Tomás Carrasco-Valenzuela, Clément Goubert, Mourdas Mohamed, as well as Nicolas Parisot, for general bioinformatic support and Marie Fablet for insights on the manuscript. This work was made possible thanks to the computing facilities of the Laboratoire de Biométrie Biologie Evolutive/Pôle Rhône-Alpes de bioinformatique (CC LBBE/PRABI), and the Symbiotron platform belonging to the FR3728 BioEEnVis. The authors used AI-based language tools to assist in improving the clarity and correctness of the writing. All content was subsequently reviewed, revised, and approved by the authors, who take full responsibility for the integrity of the work.

## References

1. Alonge, Michael, Ludivine Lebeigle, Melanie Kirsche, et al. 2022. ‘Automated Assembly Scaffolding Using RagTag Elevates a New Tomato System for High-Throughput Genome Editing’. Genome Biology 23 (1): 258. 10.1186/s13059-022-02823-7.

2. Alruiz, José M., Ignacio Peralta-Maraver, Francisco Bozinovic, Mauro Santos, and Enrico L. Rezende. 2023. ‘Temperature Adaptation and Its Impact on the Shape of Performance Curves in Drosophila Populations’. Proceedings of the Royal Society B: Biological Sciences 290 (1998): 20230507. 10.1098/rspb.2023.0507.

3. Arking, Robert, Allan G. Force, Steven P. Dudas, Steven Buck, and George T. Baker. 1996. ‘Factors Contributing to the Plasticity of the Extended Longevity Phenotypes of Drosophila’. Experimental Gerontology 31 (6): 623–43. 10.1016/S0531-5565(96)00096-4.

4. Asahina, Kenta. 2018. ‘Sex Differences in Drosophila Behavior: Qualitative and Quantitative Dimorphism’. Current Opinion in Physiology, Sex Differences, vol. 6 (December): 35–45. 10.1016/j.cophys.2018.04.004.

5. Barckmann, Bridlin, Marianne El-Barouk, Alain Pélisson, et al. 2018. ‘The Somatic piRNA Pathway Controls Germline Transposition over Generations’. Nucleic Acids Research 46 (18): 9524–36. 10.1093/nar/gky761.

6. Bates, Douglas, Martin Maechler, Ben Bolker [aut, et al. 2025. Lme4: Linear Mixed-Effects Models Using ‘Eigen’ and S4. V. 1.1-36. Released January 11. https://cran.r-project.org/web/packages/lme4/index.html.

7. Ben Shachar, Mattan, Daniel Lüdecke, and Dominique Makowski. 2020. ‘Effectsize: Estimation of Effect Size Indices and Standardized Parameters Aims of the Package’. The Journal of Open Source Software 5 (December): 2815. 10.21105/joss.02815.

8. Berrigan, David. 1997. ‘Acclimation of Metabolic Rate in Response to Developmental Temperature in Drosophila Melanogaster’. Journal of Thermal Biology 22 (3): 213–18. 10.1016/S0306-4565(97)00015-6.

9. Bolar, Kartikeya. 2019. STAT: Interactive Document for Working with Basic Statistical Analysis. V. 0.1.0. Released April 1. https://cran.r-project.org/web/packages/STAT/index.html.

10. Brasset, E., A. R. Taddei, F. Arnaud, et al. 2006. ‘Viral Particles of the Endogenous Retrovirus ZAM from Drosophila Melanogaster Use a Pre-Existing Endosome/Exosome Pathway for Transfer to the Oocyte’. Retrovirology 3 (1): 25. 10.1186/1742-4690-3-25.

11. Brennecke, Julius, Colin D. Malone, Alexei A. Aravin, Ravi Sachidanandam, Alexander Stark, and Gregory J. Hannon. 2008. ‘An Epigenetic Role for Maternally Inherited piRNAs in Transposon Silencing’. Science 322 (5906): 1387–92. 10.1126/science.1165171.

12. Burgess, Scott C., and Dustin J. Marshall. 2014. ‘Adaptive Parental Effects: The Importance of Estimating Environmental Predictability and Offspring Fitness Appropriately’. Oikos 123 (7): 769–76. 10.1111/oik.01235.

13. Cabral-de-Mello, Diogo C., and Octavio M. Palacios-Gimenez. 2024. ‘Repetitive DNAs: The “Invisible” Regulators of Insect Adaptation and Speciation’. Current Opinion in Insect Science, November, 101295. 10.1016/j.cois.2024.101295.

14. Capy, Pierre, Giuliano Gasperi, Christian Biémont, and Claude Bazin. 2000. ‘Stress and Transposable Elements: Co-Evolution or Useful Parasites?’ Heredity 85 (2): 101–6. 10.1046/j.1365-2540.2000.00751.x.

15. Casacuberta, Elena, and Josefa González. 2013. ‘The Impact of Transposable Elements in Environmental Adaptation’. Molecular Ecology 22 (6): 1503–17. 10.1111/mec.12170.

16. Catlin, Nathan S., and Emily B. Josephs. 2022. ‘The Important Contribution of Transposable Elements to Phenotypic Variation and Evolution’. Current Opinion in Plant Biology 65 (February): 102140. 10.1016/j.pbi.2021.102140.

17. Chakraborty, Mahul, J. J. Emerson, Stuart J. Macdonald, and Anthony D. Long. 2019. ‘Structural Variants Exhibit Widespread Allelic Heterogeneity and Shape Variation in Complex Traits’. Nature Communications 10 (1): 4872. 10.1038/s41467-019-12884-1.

18. Chakraborty, Mahul, Nicholas W. VanKuren, Roy Zhao, Xinwen Zhang, Shannon Kalsow, and J. J. Emerson. 2018. ‘Hidden Genetic Variation Shapes the Structure of Functional Elements in Drosophila’. Nature Genetics 50 (1): 20–25. 10.1038/s41588-017-0010-y.

19. Chandegra, Bhakti, Jocelyn Lok Yee Tang, Haoyu Chi, and Nazif Alic. 2017. ‘Sexually Dimorphic Effects of Dietary Sugar on Lifespan, Feeding and Starvation Resistance in Drosophila’. Aging 9 (12): 2521–28. 10.18632/aging.101335.

20. Choi, Jae Young, and Yuh Chwen G. Lee. 2020. ‘Double-Edged Sword: The Evolutionary Consequences of the Epigenetic Silencing of Transposable Elements’. PLOS Genetics 16 (7): e1008872. 10.1371/journal.pgen.1008872.

21. Christensen, Rune Haubo B. (2018) 2025. Runehaubo/lmerTestR. HTML. January 17, released January 14. https://github.com/runehaubo/lmerTestR.

22. Chuong, Edward B., Nels C. Elde, and Cédric Feschotte. 2017. ‘Regulatory Activities of Transposable Elements: From Conflicts to Benefits’. Nature Reviews Genetics 18 (2): 2. 10.1038/nrg.2016.139.

23. Colwell, Robert K. 1974. ‘Predictability, Constancy, and Contingency of Periodic Phenomena’. Ecology 55 (5): 1148–53. 10.2307/1940366.

24. Costantini, David. 2024. ‘For Better or Worse: How Early Life Oxidative Stress Moulds the Phenotype’. In The Role of Organismal Oxidative Stress in the Ecology and Life-History Evolution of Animals, edited by David Costantini. Springer Nature Switzerland. 10.1007/978-3-031-65183-0_7.

25. Cranz-Mileva, Susanne, Eve Reilly, Noor Chalhoub, et al. 2024. ‘Transposon Removal Reveals Their Adaptive Fitness Contribution’. Genome Biology and Evolution 16 (2): evae010. 10.1093/gbe/evae010.

26. Danecek, Petr, James K. Bonfield, Jennifer Liddle, et al. 2021. ‘Twelve Years of SAMtools and BCFtools’. GigaScience 10 (2): giab008. 10.1093/gigascience/giab008.

27. David, J., Y. Cohet, and P. Fouillet. 1975. ‘The Variability between Individuals as a Measure of Senescence: A Study of the Number of Eggs Laid and the Percentage of Hatched Eggs in the Case of Drosophila Melanogaster’. Experimental Gerontology 10 (1): 17–25. 10.1016/0531-5565(75)90011-X.

28. David, Jean R., and Pierre Capy. 1988. ‘Genetic Variation of Drosophila Melanogaster Natural Populations’. Trends in Genetics 4 (4): 106–11. 10.1016/0168-9525(88)90098-4.

29. De Coster, Wouter, and Rosa Rademakers. 2023. ‘NanoPack2: Population-Scale Evaluation of Long-Read Sequencing Data’. Bioinformatics 39 (5): btad311. 10.1093/bioinformatics/btad311.

30. Drongitis, Denise, Francesco Aniello, Laura Fucci, and Aldo Donizetti. 2019. ‘Roles of Transposable Elements in the Different Layers of Gene Expression Regulation’. International Journal of Molecular Sciences 20 (22): 22. 10.3390/ijms20225755.

31. Dubin, Manu J., Ortrun Mittelsten Scheid, and Claude Becker. 2018. ‘Transposons: A Blessing Curse’. Current Opinion in Plant Biology, 42 Genome studies and molecular genetics 2018, vol. 42 (April): 23–29. 10.1016/j.pbi.2018.01.003.

32. Dunkler, Daniela, Maria Haller, Rainer Oberbauer, and Georg Heinze. 2020. ‘To Test or to Estimate? P-Values versus Effect Sizes’. Transplant International 33 (1): 50–55. 10.1111/tri.13535.

33. Flatt, Thomas. 2020. ‘Life-History Evolution and the Genetics of Fitness Components in Drosophila Melanogaster’. Genetics 214 (1): 3–48. 10.1534/genetics.119.300160.

34. Fueyo, Raquel, Julius Judd, Cedric Feschotte, and Joanna Wysocka. 2022. ‘Roles of Transposable Elements in the Regulation of Mammalian Transcription’. Nature Reviews Molecular Cell Biology 23 (7): 481–97. 10.1038/s41580-022-00457-y.

35. Galbraith, James D., and Alexander Hayward. 2023. ‘The Influence of Transposable Elements on Animal Colouration’. Trends in Genetics 0 (0). 10.1016/j.tig.2023.04.005.

36. Gasser, M., M. Kaiser, D. Berrigan, and S. C. Stearns. 2000. ‘LIFE-HISTORY CORRELATES OF EVOLUTION UNDER HIGH AND LOW ADULT MORTALITY’. Evolution 54 (4): 1260–72. 10.1111/j.0014-3820.2000.tb00559.x.

37. Gautier, Mathieu. 2015. ‘Genome-Wide Scan for Adaptive Divergence and Association with Population-Specific Covariates’. Genetics 201 (4): 1555–79. 10.1534/genetics.115.181453.

38. Gebrie, Alemu. 2023. ‘Transposable Elements as Essential Elements in the Control of Gene Expression’. Mobile DNA 14 (August): 9. 10.1186/s13100-023-00297-3.

39. Gilbert, Clément, Jean Peccoud, and Richard Cordaux. 2021. ‘Transposable Elements and the Evolution of Insects’. Annual Review of Entomology 66 (Volume 66, 2021): 355–72. 10.1146/annurev-ento-070720-074650.

40. Groza, Cristian, Xun Chen, Travis J. Wheeler, Guillaume Bourque, and Clément Goubert. 2024. ‘A Unified Framework to Analyze Transposable Element Insertion Polymorphisms Using Graph Genomes’. Nature Communications 15 (1): 8915. 10.1038/s41467-024-53294-2.

41. Guio, Lain, and Josefa González. 2019. ‘New Insights on the Evolution of Genome Content: Population Dynamics of Transposable Elements in Flies and Humans’. In Evolutionary Genomics: Statistical and Computational Methods, edited by Maria Anisimova. Springer. 10.1007/978-1-4939-9074-0_16.

42. Guio, Lain, Cristina Vieira, and Josefa González. 2018. ‘Stress Affects the Epigenetic Marks Added by Natural Transposable Element Insertions in Drosophila Melanogaster’. Scientific Reports 8 (1): 1. 10.1038/s41598-018-30491-w.

43. Guo, Ya-Long, Jesper S. Bechsgaard, Tanja Slotte, et al. 2009. ‘Recent Speciation of Capsella Rubella from Capsella Grandiflora, Associated with Loss of Self-Incompatibility and an Extreme Bottleneck’. Proceedings of the National Academy of Sciences 106 (13): 5246–51. 10.1073/pnas.0808012106.

44. Gutzeit, Herwig O., and Roswitha Koppa. 1982. ‘Time-Lapse Film Analysis of Cytoplasmic Streaming during Late Oogenesis of Drosophila’. Development 67 (1): 101–11. 10.1242/dev.67.1.101.

45. Hall, Michael B. 2022. ‘Rasusa: Randomly Subsample Sequencing Reads to a Specified Coverage’. Journal of Open Source Software 7 (69): 3941. 10.21105/joss.03941.

46. Hayward, April, and James F. Gillooly. 2011. ‘The Cost of Sex: Quantifying Energetic Investment in Gamete Production by Males and Females’. PLOS ONE 6 (1): e16557. 10.1371/journal.pone.0016557.

47. Hirsch, Cory D., and Nathan M. Springer. 2017. ‘Transposable Element Influences on Gene Expression in Plants’. Biochimica et Biophysica Acta (BBA) - Gene Regulatory Mechanisms, Plant Gene Regulatory Mechanisms and Networks, vol. 1860 (1): 157–65. 10.1016/j.bbagrm.2016.05.010.

48. Horváth, Vivien, Miriam Merenciano, and Josefa González. 2017. ‘Revisiting the Relationship between Transposable Elements and the Eukaryotic Stress Response’. Trends in Genetics 33 (11): 832–41. 10.1016/j.tig.2017.08.007.

49. Huang, Yuheng, and Yuh Chwen G. Lee. 2024. ‘Blessing or Curse: How the Epigenetic Resolution of Host-Transposable Element Conflicts Shapes Their Evolutionary Dynamics’. Proceedings of the Royal Society B: Biological Sciences 291 (2020): 20232775. 10.1098/rspb.2023.2775.

50. Huang, Yuheng, Harsh Shukla, and Yuh Chwen G. Lee. 2022. ‘Species-Specific Chromatin Landscape Determines How Transposable Elements Shape Genome Evolution’. eLife 11 (August): e81567. 10.7554/eLife.81567.

51. Ito, Hidetaka. 2022. ‘Environmental Stress and Transposons in Plants’. Genes & Genetic Systems 97 (4): 169–75. 10.1266/ggs.22-00045.

52. Jeffreys, Harold. 1948. Theory Of Probability. http://archive.org/details/in.ernet.dli.2015.2608.

53. Joly-Lopez, Zoé, Ewa Forczek, Emilio Vello, Douglas R. Hoen, Akiko Tomita, and Thomas E. Bureau. 2017. ‘Abiotic Stress Phenotypes Are Associated with Conserved Genes Derived from Transposable Elements’. Frontiers in Plant Science 8 (November). 10.3389/fpls.2017.02027.

54. Kassambara, Alboukadel, Marcin Kosinski, Przemyslaw Biecek, and Scheipl Fabian. 2021. Survminer: Drawing Survival Curves Using ‘Ggplot2’. V. 0.4.9. Released March 9. https://cloud.r-project.org/web/packages/survminer/index.html.

55. Khazaeli, Aziz A., Wayne Van Voorhies, and James W. Curtsinger. 2005. ‘The Relationship between Life Span and Adult Body Size Is Highly Strain-Specific in Drosophila Melanogaster’. Experimental Gerontology 40 (5): 377–85. 10.1016/j.exger.2005.02.004.

56. Kolmogorov, Mikhail, Jeffrey Yuan, Yu Lin, and Pavel A. Pevzner. 2019. ‘Assembly of Long, Error-Prone Reads Using Repeat Graphs’. Nature Biotechnology 37 (5): 540–46. 10.1038/s41587-019-0072-8.

57. Kozłowski, Jan, Marek Konarzewski, and Marcin Czarnoleski. 2020. ‘Coevolution of Body Size and Metabolic Rate in Vertebrates: A Life-History Perspective’. Biological Reviews 95 (5): 1393–417. 10.1111/brv.12615.

58. Lanciano, Sophie, and Marie Mirouze. 2018. ‘Transposable Elements: All Mobile, All Different, Some Stress Responsive, Some Adaptive?’ Current Opinion in Genetics & Development, Genome Architecture and Expression, vol. 49 (April): 106–14. 10.1016/j.gde.2018.04.002.

59. Latzel, Vít, Javier Puy, Michael Thieme, Etienne Bucher, Lars Götzenberger, and Francesco de Bello. 2023. ‘Phenotypic Diversity Influenced by a Transposable Element Increases Productivity and Resistance to Competitors in Plant Populations’. Journal of Ecology 111 (11): 2376–87. 10.1111/1365-2745.14185.

60. Leblanc, P., S. Desset, B. Dastugue, and C. Vaury. 1997. ‘Invertebrate Retroviruses: ZAM a New Candidate in D.Melanogaster’. The EMBO Journal 16 (24): 7521–31. 10.1093/emboj/16.24.7521.

61. Lefranc, Agnès, and Jørgen Bundgaard. 2000. ‘The Influence of Male and Female Body Size on Copulation Duration and Fecundity in Drosophila Melanogaster’. Hereditas 132 (3): 243–47. 10.1111/j.1601-5223.2000.00243.x.

62. Li, Heng. 2021. ‘New Strategies to Improve Minimap2 Alignment Accuracy’. Bioinformatics 37 (23): 4572–74. 10.1093/bioinformatics/btab705.

63. Lighton, John R. B., and John R. B. Lighton. 2021. Measuring Metabolic Rates: A Manual for Scientists. Oxford University Press.

64. Liu, Yan-Nan, Jian-Jun Gao, Xiao-Lin Zhuang, Dong-Dong Wu, and Yan-Bo Sun. 2025. ‘Near Complete Assembly of Drosophila Melanogaster Canton S Strain Genome’. Nature Communications, ahead of print, December 3. 10.1038/s41467-025-67031-w.

65. Long, Jacob A. 2024. ‘Jtools: Analysis and Presentation of Social Scientific Data’. Journal of Open Source Software 9 (101): 6610. 10.21105/joss.06610.

66. Mackay, Trudy F. C. 1986. ‘Transposable Element-Induced Fitness Mutations in Drosophila Melanogaster’. Genetics Research 48 (2): 77–87. 10.1017/S0016672300024794.

67. Manni, Mosè, Matthew R. Berkeley, Mathieu Seppey, Felipe A. Simão, and Evgeny M. Zdobnov. 2021. ‘BUSCO Update: Novel and Streamlined Workflows along with Broader and Deeper Phylogenetic Coverage for Scoring of Eukaryotic, Prokaryotic, and Viral Genomes’. Molecular Biology and Evolution 38 (10): 4647–54. 10.1093/molbev/msab199.

68. Marin, Pierre, Julien Genitoni, Dominique Barloy, et al. 2020. ‘Biological Invasion: The Influence of the Hidden Side of the (Epi)Genome’. Functional Ecology 34 (2): 385–400. 10.1111/1365-2435.13317.

69. Mérel, Vincent, Patricia Gibert, Inessa Buch, et al. 2021. ‘The Worldwide Invasion of Drosophila Suzukii Is Accompanied by a Large Increase of Transposable Element Load and a Small Number of Putatively Adaptive Insertions’. Molecular Biology and Evolution 38 (10): 4252–67. 10.1093/molbev/msab155.

70. Merenciano, Miriam, and Josefa González. 2023. ‘The Interplay Between Developmental Stage and Environment Underlies the Adaptive Effect of a Natural Transposable Element Insertion’. Molecular Biology and Evolution 40 (3): msad044. 10.1093/molbev/msad044.

71. Merenciano, Miriam, Anaïs Larue, Cristian Groza, Cristina Vieira, Rita Rebollo, and Clément Goubert. 2024. ‘Chapter 6 - Epigenetics and Genotypic Variation: A Transposable Elements’ Perspective’. In On Epigenetics and Evolution, edited by Carlos M. Guerrero-Bosagna. Translational Epigenetics. Academic Press. 10.1016/B978-0-443-19051-3.00006-1.

72. Mikheenko, Alla, Andrey Prjibelski, Vladislav Saveliev, Dmitry Antipov, and Alexey Gurevich. 2018. ‘Versatile Genome Assembly Evaluation with QUAST-LG’. Bioinformatics 34 (13): i142–50. 10.1093/bioinformatics/bty266.

73. Minchiotti, Gabriella, Cristina Contursi, Franco Graziani, Giuseppe Gargiulo, and Pier Paolo Di Nocera. 1994. ‘Expression of Drosophila Melanogaster F Elements in Vivo’. Molecular and General Genetics MGG 245 (2): 152–59. 10.1007/BF00283262.

74. Mohamed, Mourdas, Nguyet Thi-Minh Dang, Yuki Ogyama, et al. 2020. ‘A Transposon Story: From TE Content to TE Dynamic Invasion of Drosophila Genomes Using the Single-Molecule Sequencing Technology from Oxford Nanopore’. Cells 9 (8): 8. 10.3390/cells9081776.

75. Mohamed, Mourdas, François Sabot, Marion Varoqui, et al. 2023. ‘TrEMOLO: Accurate Transposable Element Allele Frequency Estimation Using Long-Read Sequencing Data Combining Assembly and Mapping-Based Approaches’. Genome Biology 24 (1): 63. 10.1186/s13059-023-02911-2.

76. Morgante, Fabio, Peter Sørensen, Daniel A. Sorensen, Christian Maltecca, and Trudy F. C. Mackay. 2015. ‘Genetic Architecture of Micro-Environmental Plasticity in Drosophila Melanogaster’. Scientific Reports 5 (May): 9785. 10.1038/srep09785.

77. Moschetti, Roberta, Antonio Palazzo, Patrizio Lorusso, Luigi Viggiano, and René Massimiliano Marsano. 2020. ‘“What You Need, Baby, I Got It”: Transposable Elements as Suppliers of Cis-Operating Sequences in Drosophila’. Biology 9 (2): 2. 10.3390/biology9020025.

78. Negi, Pooja, Archana N. Rai, and Penna Suprasanna. 2016. ‘Moving through the Stressed Genome: Emerging Regulatory Roles for Transposons in Plant Stress Response’. Frontiers in Plant Science 7 (October). 10.3389/fpls.2016.01448.

79. Niu, Xiao-Min, Yong-Chao Xu, Zi-Wen Li, et al. 2019. ‘Transposable Elements Drive Rapid Phenotypic Variation in Capsella Rubella’. Proceedings of the National Academy of Sciences 116 (14): 6908–13. 10.1073/pnas.1811498116.

80. Okonechnikov, Konstantin, Ana Conesa, and Fernando García-Alcalde. 2016. ‘Qualimap 2: Advanced Multi-Sample Quality Control for High-Throughput Sequencing Data’. Bioinformatics 32 (2): 292–94. 10.1093/bioinformatics/btv566.

81. Oleson, Bryndon J., Daphne Bazopoulou, and Ursula Jakob. 2021. ‘Shaping Longevity Early in Life: Developmental ROS and H3K4me3 Set the Clock’. Cell Cycle 20 (22): 2337–47. 10.1080/15384101.2021.1986317.

82. Oliveira, Daniel S., Marie Fablet, Anaïs Larue, et al. 2023. ‘ChimeraTE: A Pipeline to Detect Chimeric Transcripts Derived from Genes and Transposable Elements’. Nucleic Acids Research 51 (18): 9764–84. 10.1093/nar/gkad671.

83. Park, Ah Rume, Na Liu, Nils Neuenkirchen, Qiaozhi Guo, and Haifan Lin. 2019. ‘The Role of Maternal HP1a in Early Drosophila Embryogenesis via Regulation of Maternal Transcript Production’. Genetics 211 (1): 201–17. 10.1534/genetics.118.301704.

84. Pasyukova, E. G., S. V. Nuzhdin, T. V. Morozova, and T. F. C. Mackay. 2004. ‘Accumulation of Transposable Elements in the Genome of Drosophila Melanogaster Is Associated with a Decrease in Fitness’. Journal of Heredity 95 (4): 284–90. 10.1093/jhered/esh050.

85. Piacentini, Lucia, Laura Fanti, Valeria Specchia, et al. 2014. ‘Transposons, Environmental Changes, and Heritable Induced Phenotypic Variability’. Chromosoma 123 (4): 345–54. 10.1007/s00412-014-0464-y.

86. Pimpinelli, Sergio, and Lucia Piacentini. 2020. ‘Environmental Change and the Evolution of Genomes: Transposable Elements as Translators of Phenotypic Plasticity into Genotypic Variability’. Functional Ecology 34 (2): 428–41. 10.1111/1365-2435.13497.

87. Platt, Roy N., Michael W. Vandewege, and David A. Ray. 2018. ‘Mammalian Transposable Elements and Their Impacts on Genome Evolution’. Chromosome Research 26 (1): 25–43. 10.1007/s10577-017-9570-z.

88. Pozzi, Carlo M., Angelo Gaiti, and Alberto Spada. 2025. ‘Climate Change and Plant Genomic Plasticity’. Theoretical and Applied Genetics 138 (9): 231. 10.1007/s00122-025-05010-x.

89. Pulver, Stefan, and Jimena Berni. 2012. ‘The Fundamentals of Flying: Simple and Inexpensive Strategies for Employing Drosophila Genetics in Neuroscience Teaching Laboratories’. Journal of Undergraduate Neuroscience Education : JUNE : A Publication of FUN, Faculty for Undergraduate Neuroscience 11 (October): A139–48.

90. Quinlan, Aaron R., and Ira M. Hall. 2010. ‘BEDTools: A Flexible Suite of Utilities for Comparing Genomic Features’. Bioinformatics 26 (6): 841–42. 10.1093/bioinformatics/btq033.

91. R Core Team. 2021. R: A Language and Environment for Statistical Computing. V. 4.4.0. R Foundation for Statistical Computing, Vienna, Austria., Released. https://www.R-project.org/.

92. Rebollo, Rita, Mark T. Romanish, and Dixie L. Mager. 2012. ‘Transposable Elements: An Abundant and Natural Source of Regulatory Sequences for Host Genes’. Annual Review of Genetics 46 (1): 21–42. 10.1146/annurev-genet-110711-155621.

93. Rech, Gabriel E., Santiago Radío, Sara Guirao-Rico, et al. 2022. ‘Population-Scale Long-Read Sequencing Uncovers Transposable Elements Associated with Gene Expression Variation and Adaptive Signatures in Drosophila’. Nature Communications 13 (1): 1948. 10.1038/s41467-022-29518-8.

94. Reinhold, K. 1999. ‘Energetically Costly Behaviour and the Evolution of Resting Metabolic Rate in Insects’. Functional Ecology 13 (2): 217–24. 10.1046/j.1365-2435.1999.00300.x.

95. Rey, Olivier, Etienne Danchin, Marie Mirouze, Céline Loot, and Simon Blanchet. 2016. ‘Adaptation to Global Change: A Transposable Element–Epigenetics Perspective’. Trends in Ecology & Evolution 31 (7): 514–26. 10.1016/j.tree.2016.03.013.

96. Riddell, Eric A., Marko Mutanen, and Cameron K. Ghalambor. 2023. ‘Hydric Effects on Thermal Tolerances Influence Climate Vulnerability in a High-Latitude Beetle’. Global Change Biology 29 (18): 5184–98. 10.1111/gcb.16830.

97. Roach, Michael J., Simon A. Schmidt, and Anthony R. Borneman. 2018. ‘Purge Haplotigs: Allelic Contig Reassignment for Third-Gen Diploid Genome Assemblies’. BMC Bioinformatics 19 (1): 460. 10.1186/s12859-018-2485-7.

98. Schrader, Lukas, and Jürgen Schmitz. 2019. ‘The Impact of Transposable Elements in Adaptive Evolution’. Molecular Ecology 28 (6): 1537–49. 10.1111/mec.14794.

99. Shields, Hazel J., Annika Traa, and Jeremy M. Van Raamsdonk. 2021. ‘Beneficial and Detrimental Effects of Reactive Oxygen Species on Lifespan: A Comprehensive Review of Comparative and Experimental Studies’. Frontiers in Cell and Developmental Biology 9 (February): 628157. 10.3389/fcell.2021.628157.

100. Shumate, Alaina, and Steven L. Salzberg. 2021. ‘Liftoff: Accurate Mapping of Gene Annotations’. Bioinformatics 37 (12): 1639–43. 10.1093/bioinformatics/btaa1016.

101. Slotkin, R. Keith, and Robert Martienssen. 2007. ‘Transposable Elements and the Epigenetic Regulation of the Genome’. Nature Reviews Genetics 8 (4): 272–85. 10.1038/nrg2072.

102. Smit, A. F. A., R. Hubley, and P. Green. 2021. ‘2013–2015. RepeatMasker Open-4.0’.

103. Sullivan, Gail M., and Richard Feinn. 2012. ‘Using Effect Size—or Why the P Value Is Not Enough’. Journal of Graduate Medical Education 4 (3): 279–82. 10.4300/JGME-D-12-00156.1.

104. Sundaram, Vasavi, and Joanna Wysocka. 2020. ‘Transposable Elements as a Potent Source of Diverse Cis-Regulatory Sequences in Mammalian Genomes’. Philosophical Transactions of the Royal Society B: Biological Sciences 375 (1795): 20190347. 10.1098/rstb.2019.0347.

105. Therneau, Terry M., Thomas Lumley (original S.->R port and R. maintainer until 2009), Atkinson Elizabeth, and Crowson Cynthia. 2024. Survival: Survival Analysis. V. 3.8-3. Released December 17. https://cran.r-project.org/web/packages/survival/index.html.

106. Van Voorhies, Wayne A. 2009. ‘Metabolic Function in Drosophila Melanogaster in Response to Hypoxia and Pure Oxygen’. Journal of Experimental Biology 212 (19): 3132–41. 10.1242/jeb.031179.

107. Varoqui, Marion, Mourdas Mohamed, Bruno Mugat, et al. 2025. ‘Temporal and Spatial Niche Partitioning in a Retrotransposon Community of the Drosophila Melanogaster Genome’. Nucleic Acids Research 53 (11): gkaf516. 10.1093/nar/gkaf516.

108. Vaser, Robert, Ivan Sović, Niranjan Nagarajan, and Mile Šikić. 2017. ‘Fast and Accurate de Novo Genome Assembly from Long Uncorrected Reads’. Genome Research 27 (5): 737–46. 10.1101/gr.214270.116.

109. Vaury, C., M. C. Chaboissier, M. E. Drake, O. Lajoinie, B. Dastugue, and A. Pélisson. 1994. ‘The Doc Transposable Element in Drosophila Melanogaster and Drosophila Simulans: Genomic Distribution and Transcription’. Genetica 93 (1–3): 117–24. 10.1007/BF01435244.

110. Vermeulen, C. J., and R. Bijlsma. 2004. ‘Changes in Mortality Patterns and Temperature Dependence of Lifespan in Drosophila Melanogaster Caused by Inbreeding’. Heredity 92 (4): 275–81. 10.1038/sj.hdy.6800412.

111. Videlier, Mathieu, Howard D. Rundle, and Vincent Careau. 2019. ‘Sex-Specific Among-Individual Covariation in Locomotor Activity and Resting Metabolic Rate in Drosophila Melanogaster’. The American Naturalist 194 (6): E164–76. 10.1086/705678.

112. Videlier, Mathieu, Howard D. Rundle, and Vincent Careau. 2021. ‘Sex-specific Genetic (Co)Variances of Standard Metabolic Rate, Body Mass and Locomotor Activity in Drosophila Melanogaster’. Journal of Evolutionary Biology 34 (8): 1279–89. 10.1111/jeb.13887.

113. Weilguny, Lukas, and Robert Kofler. 2019. ‘DeviaTE: Assembly-Free Analysis and Visualization of Mobile Genetic Element Composition’. Molecular Ecology Resources 19 (5): 1346–54. 10.1111/1755-0998.13030.

114. Wells, Jonathan N., and Cédric Feschotte. 2020. ‘A Field Guide to Eukaryotic Transposable Elements’. Annual Review of Genetics 54 (November): 539–61. 10.1146/annurev-genet-040620-022145.

115. Wheeler, Matthew W., Robert M. Park, and A. John Bailer. 2006. ‘Comparing Median Lethal Concentration Values Using Confidence Interval Overlap or Ratio Tests’. Environmental Toxicology and Chemistry 25 (5): 1441–44. 10.1897/05-320r.1.

116. Whitelaw, Emma, and David I. K. Martin. 2001. ‘Retrotransposons as Epigenetic Mediators of Phenotypic Variation in Mammals’. Nature Genetics 27 (4): 4. 10.1038/86850.

117. Woodruff, R. C., J. N. Thompson, J. S. F. Barker, and H. Huai. 2000. ‘Transposable DNA Elements and Life History Traits: II. Transposition of P DNA Elements in Somatic Cells Reduces Fitness, Mating Activity, and Locomotion of Drosophila Melanogaster’. In Transposable Elements and Genome Evolution, edited by John F. McDonald. Springer Netherlands. 10.1007/978-94-011-4156-7_26.

118. Zwaan, B. J., R. Bijlsma, and R. F. Hoekstra. 1992. ‘On the Developmental Theory of Ageing. II. The Effect of Developmental Temperature on Longevity in Relation to Adult Body Size in D. Melanogaster’. Heredity 68 (2): 123–30. 10.1038/hdy.1992.19.

