## Supplementary document for "Transposable elements drive phenotypic variation and shape the response to environmental changes in *Drosophila melanogaster*"

Supplementary material ... 3
Figure S1: Newly detected LTR insertions relative to G0. ... 3
Figure S2: Newly detected LINE insertions relative to G0. ... 4
Figure S3: Newly detected DNA transposon insertions relative to G0. ... 5
Supplementary analysis: non-repeat structural variants support limited population divergence ... 6
Table S1: Summary statistics of the non-repeat SVs frequencies at the population scale. ... 7
Figure S4: Size and frequency distributions of the non-repeat structural variants (SVs). ... 8
Table S2: Ranking of the standard deviation values (SD) of female traits across populations. ... 9
Figure S5: Transposable element abundance as estimated by DeviaTE in the TE-accumulation populations ... 10
Table S3: Slope of the reaction norm of LT80 under temperature fluctuation. ... 10
Table S4: Results of a three-way analysis of variance (ANOVA) from the LMER model testing the effects of the population, temperature, as well as their interactions, on females' metabolic rate ... 11
Table S5: Results of a three-way analysis of variance (ANOVA) from the LMER model testing the effects of the population, temperature, as well as their interactions, on males' metabolic rate. ... 11
Table S6: Results of a three-way analysis of variance (ANOVA) from the LMER model of the effect of population, paraquat concentration, and their interaction on females' longevity. ... 12
Table S7: Results of a three-way analysis of variance (ANOVA) from the LMER model of the effect of population, paraquat concentration, and their interaction on males' longevity. ... 12
Table S8: Slope of the reaction norm of LT80 in response to changes in paraquat concentration ... 12
Table S9: Estimates of LT80 under variable paraquat concentrations in TE-accumulation populations. ... 13
Table S10: Manually curated candidate transposable element insertions. ... 14
Supplementary table legends: ... 15
Table S11: Genome-scan candidate transposable element (TE) insertions with manual curation and integration of female phenotypic data. ... 15
Table S12: Genome-scan candidate transposable element (TE) insertions with manual curation and integration of male phenotypic data. ... 15
Table S14: Assembly completeness and quality metrics of the genome assemblies. ... 16
Figure S6: Rarefaction curves of new TE insertions detected by TrEMOLO in comparison to G0 across the TE-accumulation populations. ... 17
Figure S7: Experimental design for the viability and hatchability assays. ... 18

### Supplementary material

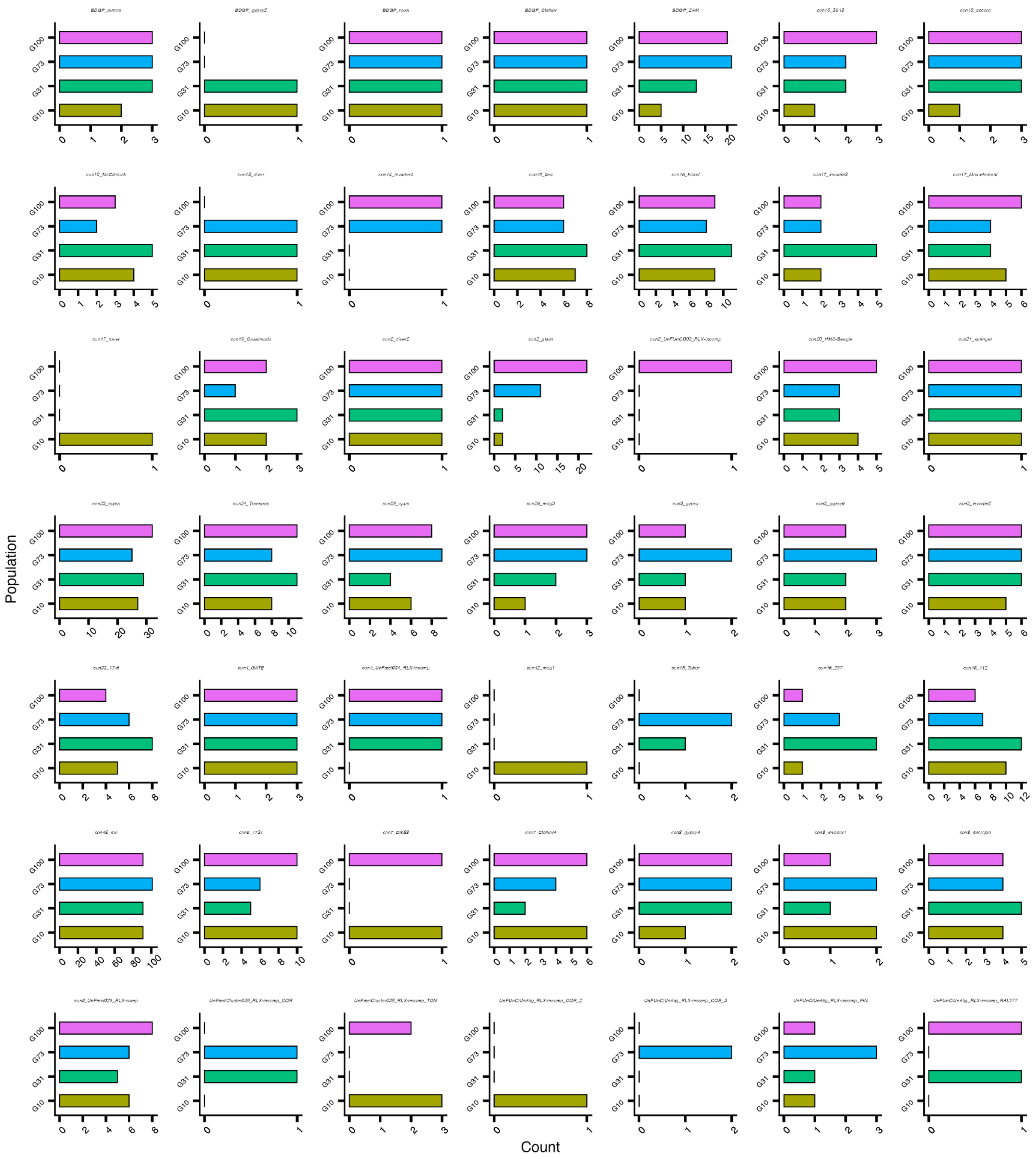

***Figure S1: Newly detected LTR insertions relative to G0.***

*Each bar represents the count of insertions identified in the sample that are absent in the G0 reference, as detected by TrEMOLO. Insertions are categorized by family according to the consensus library.*

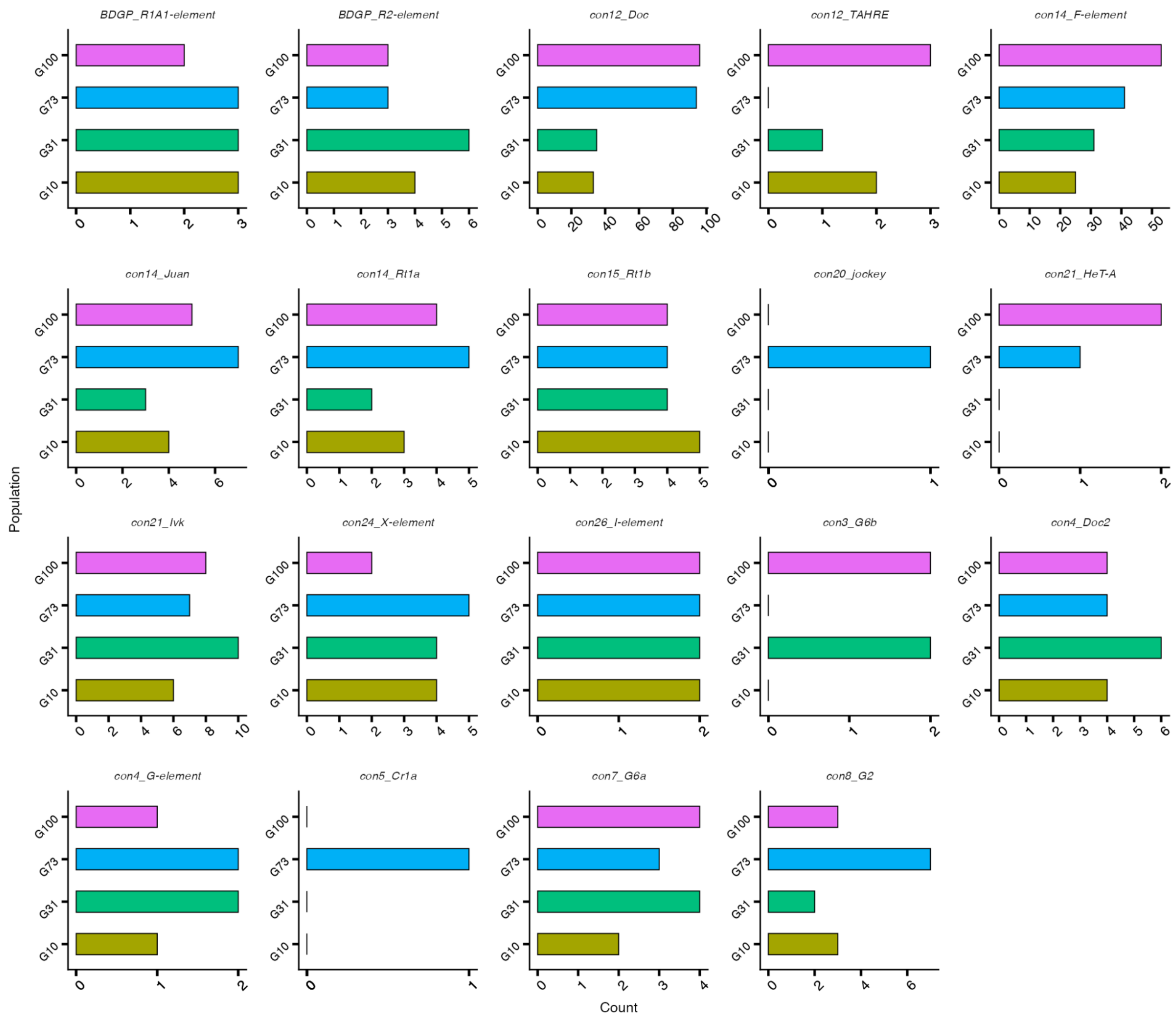

***Figure S2: Newly detected LINE insertions relative to G0.***

*Each bar represents the count of insertions identified in the sample that are absent in the G0 reference, as detected by TrEMOLO. Insertions are categorized by family according to the consensus library.*

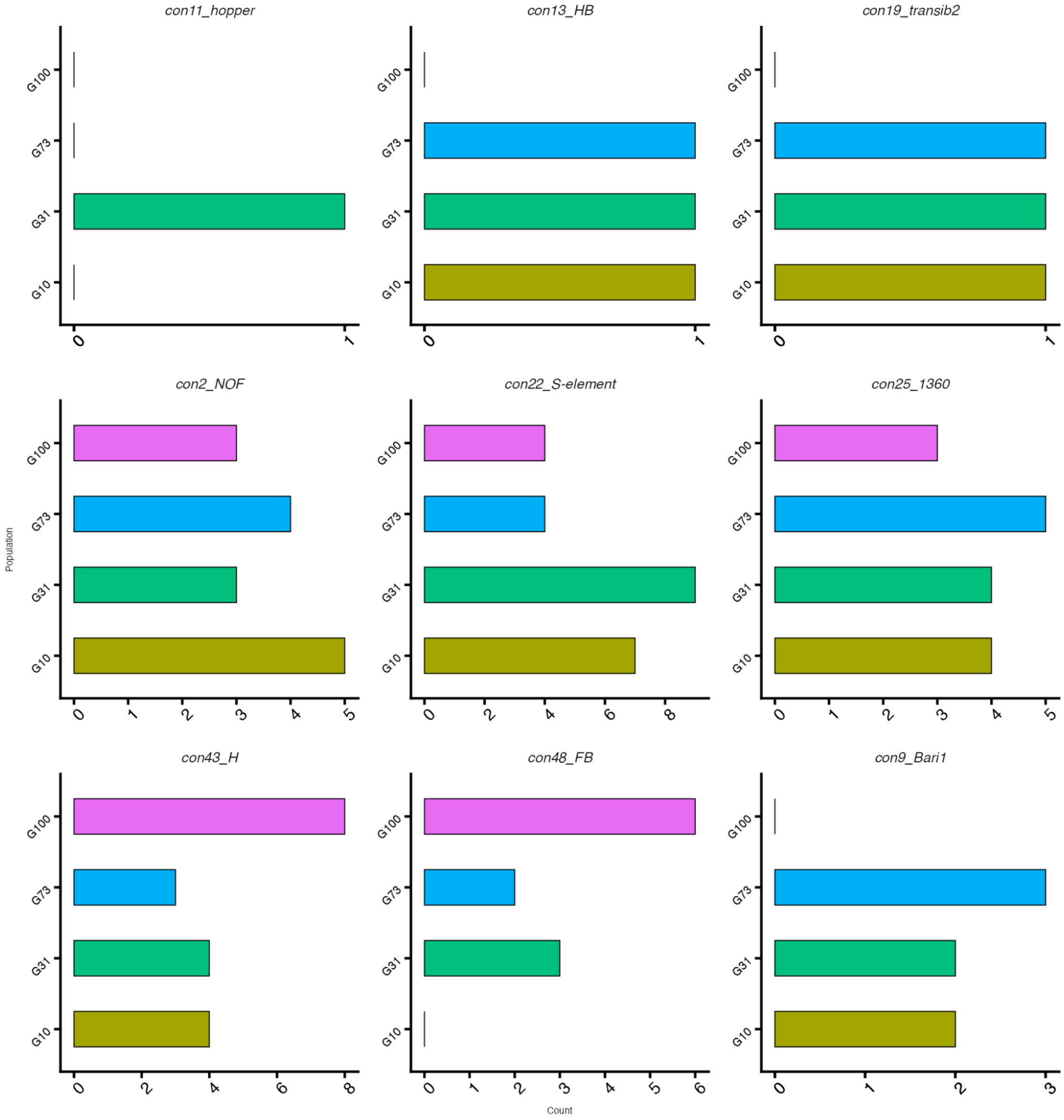

***Figure S3: Newly detected DNA transposon insertions relative to G0.***

*Each bar represents the count of insertions identified in the sample that are absent in the G0 reference, as detected by TrEMOLO. Insertions are categorized by family according to the consensus library.*

### Supplementary analysis: non-repeat structural variants support limited population divergence

Non-repeat SVs (non-TE and non-satellite SVs) were obtained by diverting the GraffiTE pipeline (Groza et al. 2024) (see Material and Methods). SV calling was performed on the genome assemblies and compared to G0, not to obtain an exhaustive list of all SVs that might be present in individual genomes, but rather to provide a general overview of SVs at the population scale. The reason for this approach is to address the putative influence of non-repeat SVs on the phenotype at the population scale. First, it is important to note that the number of deletions and insertions detected in the assemblies does not increase across generations of TE accumulation (Table S1). The number of non-repeat deletions differing from G0 is 68 in G10, and 61, 66, and 57 in G31, G73, and G100, respectively (Table S1). For insertions, G10 carries 45 non-repeat insertions not detected in the G0 reference genome, while G31, G73, and G100 carry 59, 47, and 54, respectively (Table S1). While deletions appear to be population-specific, insertions show a higher degree of shared presence, with 13 insertions common to all four populations (Figure S4).

The size distribution of these indels reveals that most fall in the range of 100-500 bp, with an average of 85% of all indels within this size range. Additionally, no significant differences in size distribution were observed, further supporting that non-repeat SVs contribute only marginally to the genotypic divergence of TE-accumulation populations with respect to SV size (p-value = 0.6495) (Figure S4A). The frequencies of each SV called from the genome assemblies were inferred by aligning reads to the SVs (see Material and Methods). Non-repeat SV frequencies did not differ significantly between populations (Kruskal-Wallis p-value = 0.08114) (Figure S4B). The average deletion frequency is approximately 55%, suggesting that most deletions are present in a heterozygous state, while the average insertion frequency is around 78%.

Notably, the amount of non-repeat SVs detected between the assemblies is drastically lower than those reported in the literature. Indeed, while we report an average of, respectively, 63 and 51.25 deletions and insertions, in one assembly compared to G0 (Table S1), the comparison of a reference-quality genome of strain A4 to *dm6* uncovered 404 non-TE indels, of which 223 were insertions and the remainder deletions (Chakraborty et al. 2018). In a study comparing 14 different *D. melanogaster* strains of reference-quality genomes, the same authors detected 4,347 non-TE indels larger than 100 bp (Chakraborty et al. 2019). On average, each strain comparison to the ISO-1 strain revealed 694 non-TE indels, ranging from 584 to 916, differentiating the non-TE SVs (Chakraborty et al. 2019). More recently, a telomere-to-telomere comparison of ISO-1 and Canton S assemblies, using a combined approach with Sniffles, SIM, PBSV, and CuteSV for SV calling, detected 7,989 SVs, of which 2,862 were insertions and

3,208 were deletions (Liu et al. 2025). Only 1% of the detected SVs belonged to repetitive sequences (Liu et al. 2025). Massouras and collaborators suggested that *D. melanogaster* carries a high density of molecular polymorphisms, much greater than that observed in humans and mice (Massouras et al. 2012).

Importantly, although these non-repeat SVs were filtered for assembly alignment discordances using SVIM-asm (as implemented in GraffiTE), the assemblies were generated at different sequencing depths, making it likely that some detected SVs are artifacts rather than true biological divergence. Altogether, these results support that non-repeat SVs in the TE-accumulation populations, in relation to G0, are far fewer than expected between two *D. melanogaster* assemblies and therefore might not significantly contribute to their genetic diversity.

| Population | Type | Number | Frequency mean | Frequency median | Standard deviation | Minimum frequency |
|---|---|---|---|---|---|---|
| **G10** | **DEL** | 68 | 49.93 | 52.01 | 32.97 | 1.67 |
| | **INS** | 45 | 81.40 | 92.31 | 23.82 | 12.88 |
| **G31** | **DEL** | 61 | 62.92 | 67.59 | 33.42 | 2.50 |
| | **INS** | 59 | 81.98 | 95.74 | 25.49 | 0.12 |
| **G73** | **DEL** | 66 | 51.90 | 55.43 | 33.42 | 0.26 |
| | **INS** | 47 | 78.46 | 88.37 | 25.83 | 3.45 |
| **G100** | **DEL** | 57 | 57.36 | 54.92 | 29.09 | 2.08 |
| | **INS** | 54 | 71.74 | 91.74 | 33.27 | 1.96 |

***Table S1: Summary statistics of the non-repeat SVs frequencies at the population scale.***

*Non-repeat SVs absent from G0 were obtained with GraffiTE assembly pangenomic approach. The non-repeat SVs frequencies were computed for each population. The table reports the mean, median, standard deviation, and the minimum and maximum frequency of TE insertions for each population. "DEL": deletions and "INS": insertions.*

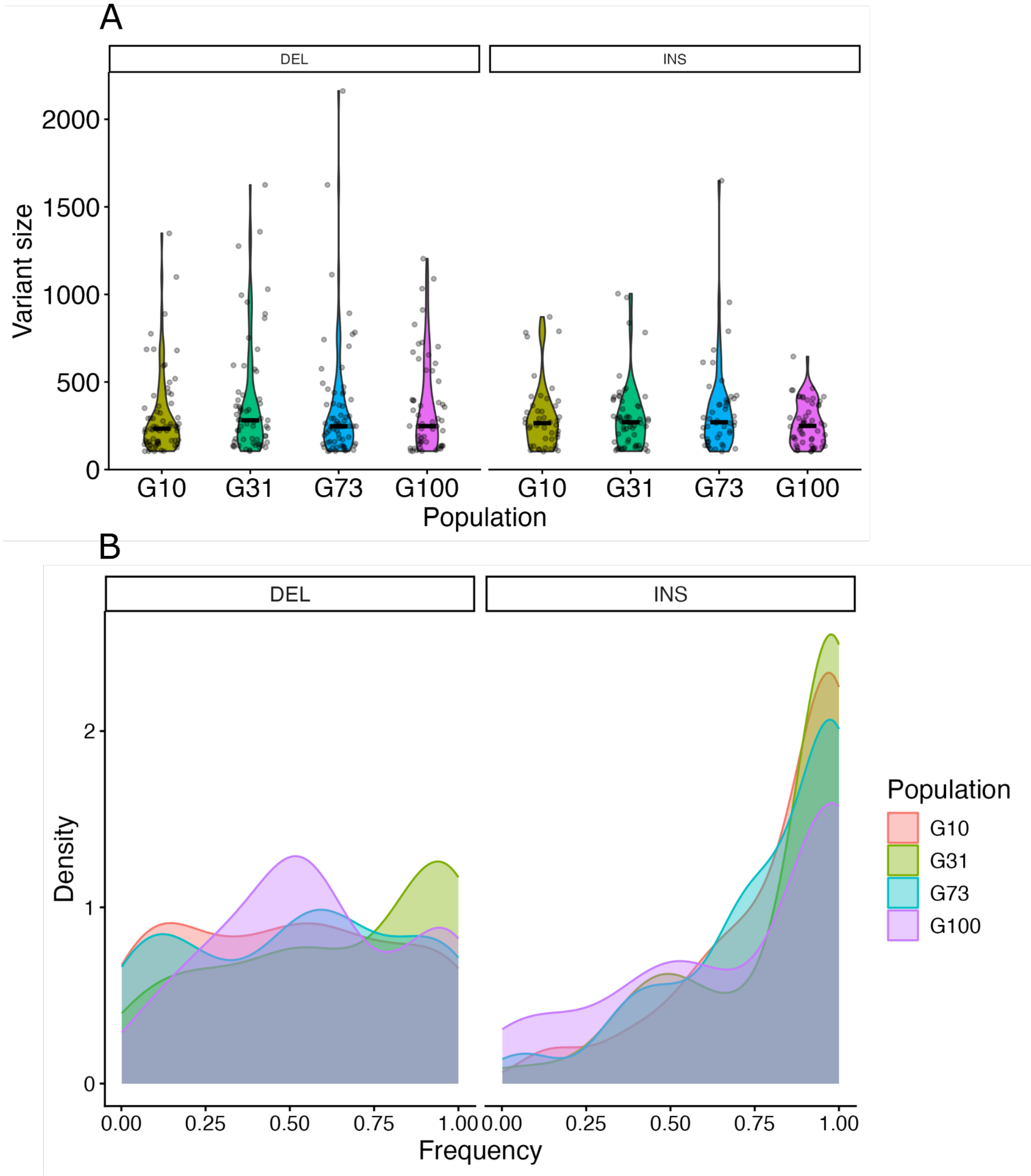

***Figure S4: Size and frequency distributions of the non-repeat structural variants (SVs).***

A) *Size distribution of the SVs, displayed as violin plots for the deletions ("DEL") on the left panel and insertions ("INS") on the right. There are no differences between the populations regarding the size of the SVs. B) Frequency distribution of the SVs displayed as a density plot for the deletions on the left and insertions on the right. There are no differences between the populations regarding the frequencies of the SVs.*

| Trait | G0 | G10 | G31 | G73 | G100 |
|---|---|---|---|---|---|
| **LT80_18** | 4 | 5 | 1 | 3 | 2 |
| **LT80_22** | 3 | 1 | 4 | 2 | 5 |
| **VCO2_18** | 5 | 4 | 3 | 1 | 2 |
| **VCO2_25** | 1 | 4 | 5 | 3 | 2 |
| **VCO2_30** | 3 | 5 | 1 | 4 | 2 |
| **Paraquat_15mM** | 2 | 3 | 1 | 5 | 4 |
| **Paraquat_25mM** | 4 | 3 | 5 | 1 | 2 |
| **Climbing** | 4 | 5 | 3 | 2 | 1 |
| **Viability** | 3 | 4 | 5 | 1 | 2 |
| **Development time** | 4 | 5 | 3 | 1 | 2 |
| **Hatchability** | 3 | 5 | 1 | 2 | 4 |
| **Dry weight** | 4 | 3 | 2 | 1 | 5 |
| **CTmax** | 5 | 4 | 1 | 2 | 3 |
| **Diet_Complete** | 5 | 3 | 2 | 1 | 4 |
| **Diet_Starvation** | 4 | 1 | 5 | 2 | 3 |
| **Diet_-Sugar** | 5 | 4 | 2 | 3 | 1 |
| **Diet_-Yeast** | 3 | 4 | 5 | 1 | 2 |

***Table S2: Ranking of the standard deviation values (SD) of female traits across populations.***

*The ranks for the 17 measured traits are reported for females in five populations. A rank of 1 corresponds to the highest SD for a given trait, indicating the greatest variability among individuals.*

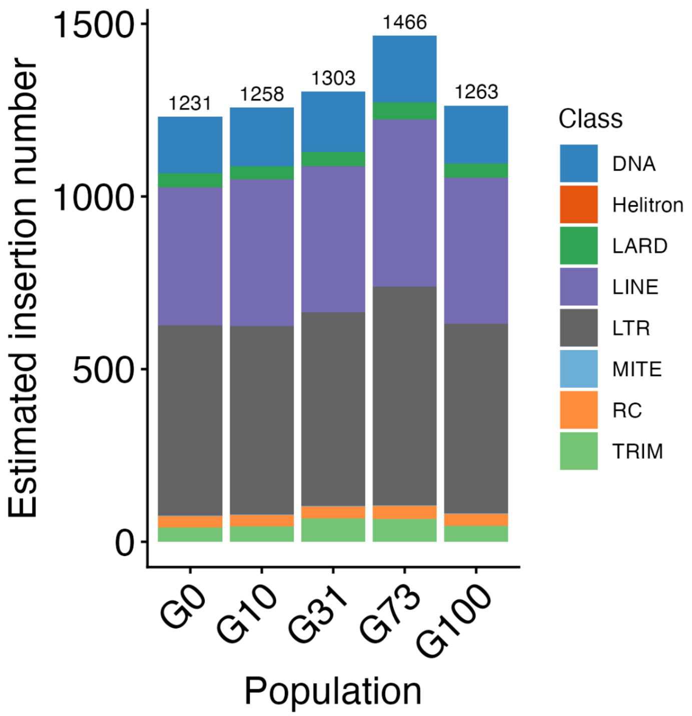

***Figure S5: Transposable element abundance as estimated by DeviaTE in the TE-accumulation populations.***

*DeviaTE enables the estimation of the overall TE quantification independent of a reference genome. In the bar plot, each colour represents a different TE subclass, and the total TE abundance estimate for each population is shown at the top of the corresponding bar.*

| Population | Sex | Slope | Average |
|---|---|---|---|
| G0 | F | -9.325964 | |
| G10 | F | -1.526493 | |
| G31 | F | -12.896336 | -7.8085056 |
| G73 | F | -5.52649 | |
| G100 | F | -9.767245 | |
| G0 | M | -4.154817 | |
| G10 | M | -4.234348 | |
| G31 | M | -2.992948 | -3.4593196 |
| G73 | M | -1.494491 | |
| G100 | M | -4.419994 | |

***Table S3: Slope of the reaction norm of LT80 under temperature fluctuation.***

*The slopes were calculated with a linear model of LT80 versus the temperature. Separate slopes are reported for each sex (F: females, M: males) for each population, along with the average slope per sex across the populations.*

| Term | Sum Sq | Mean Sq | F value | Pr(>F) |
|---|---|---|---|---|
| Temp | 1.50672 | 0.75336 | 137.99419 | 0.00000 |
| Population | 0.12215 | 0.03054 | 5.59580 | 0.00123 |
| log(DW_Fly) | 0.09698 | 0.09698 | 17.77512 | 0.00016 |
| log(Act_Fly + 1) | 0.00826 | 0.00826 | 1.51433 | 0.22450 |
| Temp:Population | 0.04442 | 0.00555 | 1.01753 | 0.43963 |

***Table S4: Results of a three-way analysis of variance (ANOVA) from the LMER model testing the effects of the population, temperature, as well as their interactions, on females' metabolic rate.***

*Term indicates the main effects or interaction terms included in the model. The table reports the sum of squares (Sum Sq), mean squares (Mean Sq), F-statistic values (F-value), and associated probabilities (Pr(>F)). Dry weight (DW) and activity (Act) were included as covariates in the model. All data were log-transformed prior to analysis. Temp: temperature, DW: dry weight, Act: activity.*

| Term | Sum Sq | Mean Sq | F value | Pr(>F) |
|---|---|---|---|---|
| Temp | 1.75513 | 0.87756 | 95.63844 | 0.00000 |
| Population | 0.16860 | 0.04215 | 4.60432 | 0.00368 |
| log(DW_Fly) | 0.01375 | 0.01375 | 1.50250 | 0.22630 |
| log(Act_Fly + 1) | 0.07339 | 0.07339 | 8.01722 | 0.00704 |
| Temp:Population | 0.12220 | 0.01528 | 1.66850 | 0.13581 |

***Table S5: Results of a three-way analysis of variance (ANOVA) from the LMER model testing the effects of the population, temperature, as well as their interactions, on males' metabolic rate.***

*Term indicates the main effects or interaction terms included in the model. The table reports the sum of squares (Sum Sq), mean squares (Mean Sq), F-statistic values (F-value), and associated probabilities (Pr(>F)). Dry weight (DW) and activity (Act) were included as covariates in the model. All data were log-transformed prior to analysis. Temp: temperature, DW: dry weight, Act: activity.*

| Term | Sum Sq | Mean Sq | F value | Pr(>F) |
|---|---|---|---|---|
| Population | 134990.07997 | 33747.51999 | 4.69642 | 0.00093 |
| Concentration | 955136.40284 | 955136.40284 | 132.92009 | 0.00000 |
| Population:Concentration | 141569.48155 | 35392.37039 | 4.92532 | 0.00062 |

**Table S6: Results of a three-way analysis of variance (ANOVA) from the LMER model of the effect of population, paraquat concentration, and their interaction on females' longevity.**

*Term indicates the main effects or interaction terms included in the model. The table reports the sum of squares (Sum Sq), mean squares (Mean Sq), F-statistic values (F-value), and associated probabilities (Pr(>F)). Dry weight (DW) and activity (Act) were included as covariates in the model.*

| Term | Sum Sq | Mean Sq | F value | Pr(>F) |
|---|---|---|---|---|
| Population | 344220.36261 | 86055.09065 | 33.86607 | 0.00000 |
| Concentration | 71343.98476 | 71343.98476 | 28.07667 | 0.00000 |
| Population:Concentration | 61832.96141 | 15458.24035 | 6.08343 | 0.00008 |

**Table S7: Results of a three-way analysis of variance (ANOVA) from the LMER model of the effect of population, paraquat concentration, and their interaction on males' longevity.**

*Term indicates the main effects or interaction terms included in the model. The table reports the sum of squares (Sum Sq), mean squares (Mean Sq), F-statistic values (F-value), and associated probabilities (Pr(>F)). Dry weight (DW) and activity (Act) were included as covariates in the model.*

| Population | Sex | Slope | Average |
|---|---|---|---|
| G0 | F | -8.8393279 | -12.1881913 |
| G10 | F | -16.8011068 | |
| G31 | F | -17.599134 | |
| G73 | F | -5.6830703 | |
| G100 | F | -12.0183175 | |
| G0 | M | -4.384834 | -2.92268918 |
| G10 | M | -1.9949077 | |
| G31 | M | 0.5274522 | |
| G73 | M | -5.9973389 | |
| G100 | M | -2.7638175 | |

**Table S8: Slope of the reaction norm of LT80 in response to changes in paraquat concentration.**

*The slopes were calculated with a linear model of LT80 versus the concentration. Separate slopes are reported for each sex (F: females, M: males) for each population, along with the average slope per sex across the populations.*

| Population | Sex | Concentration | LT80 | LCL | UCL |
|---|---|---|---|---|---|
| G0 | F | 25mM | 86.0104075 | NA | NA |
| | | 15mM | 174.403687 | 163.660409 | 186.569396 |
| | M | 25mM | 100.789266 | 94.1138279 | 108.906886 |
| | | 15mM | 144.637605 | 138.920145 | 150.928207 |
| G10 | F | 25mM | 69.6729217 | 66.9074091 | 72.7663758 |
| | | 15mM | 237.683989 | 227.490759 | 248.905344 |
| | M | 25mM | 59.9235176 | NA | NA |
| | | 15mM | 79.8725942 | 70.5686047 | 92.8409987 |
| G31 | F | 25mM | 93.2764749 | NA | NA |
| | | 15mM | 269.267815 | 256.88378 | 282.973953 |
| | M | 25mM | 103.326411 | 98.1128562 | 109.347036 |
| | | 15mM | 98.0518891 | 80.0546353 | 129.143436 |
| G73 | F | 25mM | 109.201941 | 99.4769627 | 121.123447 |
| | | 15mM | 166.032644 | 157.769222 | 175.161052 |
| | M | 25mM | 98.9805433 | 95.1878346 | 103.211389 |
| | | 15mM | 158.953932 | 153.632469 | 164.744728 |
| G100 | F | 25mM | 103.306844 | 95.9727987 | 112.053601 |
| | | 15mM | 223.490019 | 213.923413 | 233.908189 |
| | M | 25mM | 126.63997 | 121.390001 | 132.504481 |
| | | 15mM | 154.278145 | 148.570153 | 160.553743 |

***Table S9: Estimates of LT80 under variable paraquat concentrations in TE-accumulation populations.***

*The table reports the estimated time until 80% mortality is reached (LT80) along with upper and lower confidence intervals, calculated using the LT_probit function (see Material and Methods). Estimates are shown separately for females (F) and males (M) in each population. NA: not applicable.*

| TE_ID | Strand | TSD | pident | psize_TE | SIZE_TE | NEW_POS | FREQ | FREQ_WITH_CLIPPED | SV_SIZE | G0 annotation |
|---|---|---|---|---|---|---|---|---|---|---|
| con19_Quasimodo#LTR/Gypsy | - | TATATATAT | 97.346 | 98.02 | 7379 | 18047875 | 60 | 90.9091 | 7443 | 5-HT1A |
| con48_roo#LTR/Pao | + | CTCACT | 98.527 | 95.94 | 9088 | 11594593 | 80 | 97.8723 | 9161 | Ptp10D |
| con2_NOF#DNA/MULE-NOF | - | ATTC | 97.723 | 139.86 | 7385 | 14182869 | 75 | 97.7273 | 9392 | CG32599 |
| con12_Doc#LINE/Jockey | + | AGAA | 98.614 | 98.68 | 4737 | 875331 | 100 | 100 | 4788 | Myo81F |
| con12_Doc#LINE/Jockey | - | ATCAATAAA | 98.678 | 98.77 | 4725 | 939437 | 90.9091 | 97.2973 | 4795 | Myo81F |
| con12_Doc#LINE/Jockey | + | GAACTGG | 98.844 | 98.6 | 4717 | 219916 | 80 | 95.7447 | 4795 | CG3777 |
| con19_Quasimodo#LTR/Gypsy | - | ATTTTTTTT | 97.558 | 98.06 | 7395 | 12750052 | 62.5 | 92.5 | 7449 | Pde9 |
| con48_roo#LTR/Pao | + | CTAAG | 98.799 | 95.79 | 9070 | 4169725 | 80 | 97.4359 | 9136 | CG15570 |
| con14_F-element#LINE/Jockey | + | TGGCC | 98.418 | 99.28 | 4706 | 17674475 | 71.4286 | 90.6977 | 4773 | lncRNA:CR43802 |
| BDGP_ZAM#LTR/Gypsy | + | CGCG | 98.76 | 100.08 | 8442 | 21186350 | 66.6667 | 92.1569 | 8540 | CG34377 |
| con2_gtwin#LTR/Gypsy | + | AGAA | 99.057 | 99.8 | 7424 | 20902907 | 69.2308 | 93.75 | 7487 | pip |
| con48_roo#LTR/Pao\|sniffles.INS.22216 | - | GGCATTT | 98.514 | 96.09 | 9122 | 27577710 | 64.2857 | 90.1961 | 9164 | sima |

***Table S10: Manually curated candidate transposable element insertions.***

*These candidate TE insertions were selected based on their frequency (≤ 60%), their genomic location, and their insertion site according to the G0-+ gene annotation. The table reports the percent identity of each insertion with the TE consensus sequence (pident), the percentage of its size relative to the TE in the database (psize_TE), the estimated TE size (SIZE_TE), and the corrected insertion position accounting for the target site duplication (TSD) (NEW_POS). Additional information includes the insertion frequency supported by clipped reads (FREQ_WITH_CLIPPED), the corresponding gene annotation from the G0 reference, and the size of the structural variant (SV) (SV_SIZE), which can exceed the TE size.*

**Supplementary table legends:**

***Table S11: Genome-scan candidate transposable element (TE) insertions with manual curation and integration of female phenotypic data.***

*The first tab contains the allelic frequencies of the markers (excluding the sex chromosomes, and based on the number of collected embryos for sequencing, 2n = 400) used for both the female and male analyses. The second tab comprises the means and LT80 of the 17 phenotypic measurements included in the GWAS. The subsequent five spreadsheets contain the outputs of the five Baypass iterations, followed by the calculation of the median of the five BF factors (tab 7). Finally, the last tab displays the extended manual curation of the candidate insertions, accompanied by the phenotypic associations detected by Baypass, their corresponding frequencies in the TE-accumulation populations, the TE identity, and the genomic location based on G0 annotation. In red are shown the major determinants of manual curation.*

***Table S12: Genome-scan candidate transposable element (TE) insertions with manual curation and integration of male phenotypic data.***

*The first tab comprises the means and LT80 of the 16 phenotypic measurements included in the GWAS. The subsequent five spreadsheets contain the outputs of the five Baypass iterations, followed by the calculation of the median of the five BF factors (tab 6).*

***Table S13: Genome-scan of the non-repeat structural variants (SVs) for the female phenotypic data.***

*The first tab contains the allelic frequencies of the markers (excluding the sex chromosomes, and based on the number of collected embryos for sequencing, 2n = 400). The subsequent five spreadsheets contain the outputs of the five Baypass iterations, followed by the calculation of the median of the five BF factor.*

| | Complete (%) | Single Copy (%) | Duplicated (%) | Fragmented (%) | Missing (%) | Total length | N° of contigs | N50 | GC % | Coverage (mapped read) |
|---|---|---|---|---|---|---|---|---|---|---|
| G0 | 99.70 | 99.20 | 0.50 | 0.10 | 0.20 | 134182872 | 256 | 24220491 | 42.05 | 53.08 |
| G0-F100 | 99.80 | 99.20 | 0.50 | 0.10 | 0.10 | 142523898 | 77 | 25540886 | 42.14 | 107.80 |
| G10 | 99.6 | 99.1 | 0.5 | 0.2 | 0.2 | 133065001 | 206 | 24311201 | 42.11 | 65.43 |
| G31 | 99.7 | 99.2 | 0.5 | 0.1 | 0.2 | 133155141 | 155 | 24494539 | 42.10 | 38.71 |
| G73 | 99.7 | 99 | 0.7 | 0.1 | 0.2 | 133549986 | 197 | 24454906 | 42.10 | 42.24 |
| G100 | 99.6 | 99.1 | 0.5 | 0.2 | 0.2 | 132914130 | 209 | 24924766 | 42.12 | 47.45 |

***Table S14: Assembly completeness and quality metrics of the genome assemblies.***

*Assembly completeness was assessed using BUSCO, reporting percentages of complete, single-copy, duplicated, fragmented, and missing genes. Assembly contiguity and quality are summarized with the number of scaffolds and contigs, total assembly length and percentage of gaps. The grey boxes indicate the absence of metrics. The contig count, N50 (length in basepairs of the shortest contig at 50% of the total assembly length), and GC content were assessed with Quast. Coverage corresponds to the average coverage as assessed by cramino on mapping read against Dm6 reference genome. Results are shown for the G0 assembly from this study and G0-F100 assembly from* (Mohamed et al. 2020) *(available at ENA: ERP122844, and the assemblies for the TE-accumulation populations.*

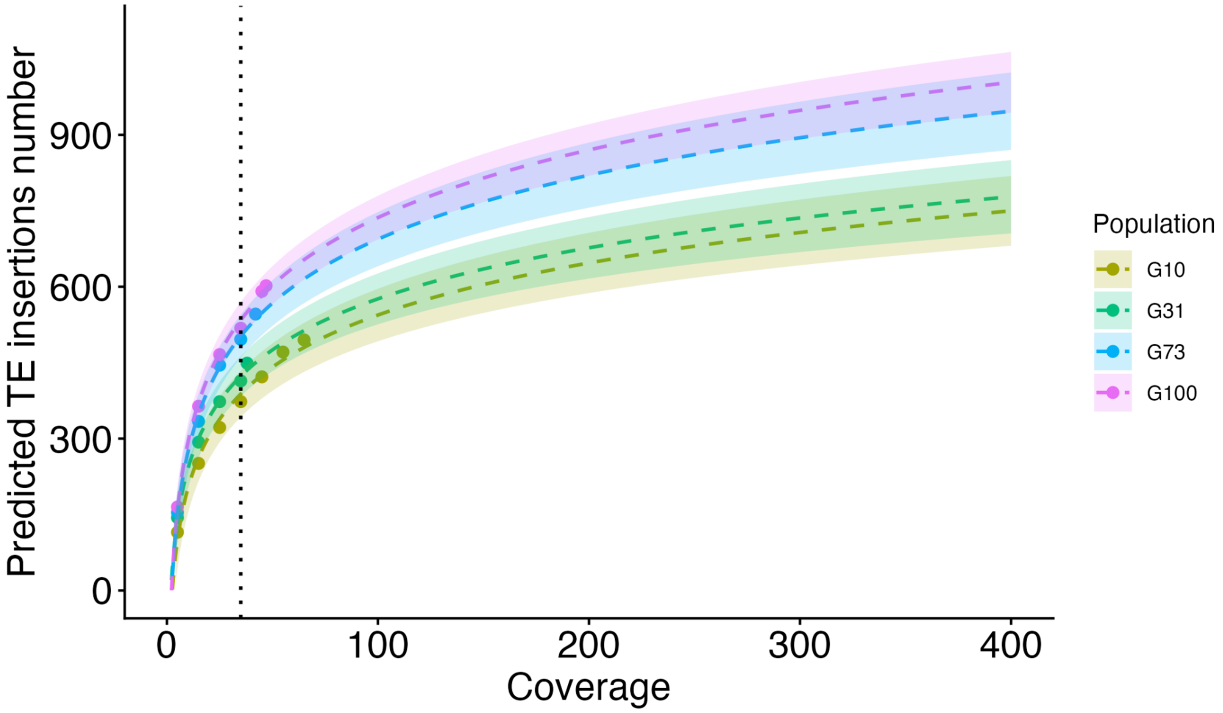

***Figure S6: Rarefaction curves of new TE insertions detected by TrEMOLO in comparison to G0 across the TE-accumulation populations.***

*The number of predicted TE insertions is plotted as a function of sequencing coverage using a logarithmic prediction function. Each dashed line represents the fitted rarefaction curve for one population, with shaded areas indicating confidence intervals. Dots correspond to observed values. The vertical dotted lines indicate selected coverage thresholds, namely 35 X. The left panel provides a zoomed-in view of the curves at lower coverage values, corresponding to the initial region of the right panel.*

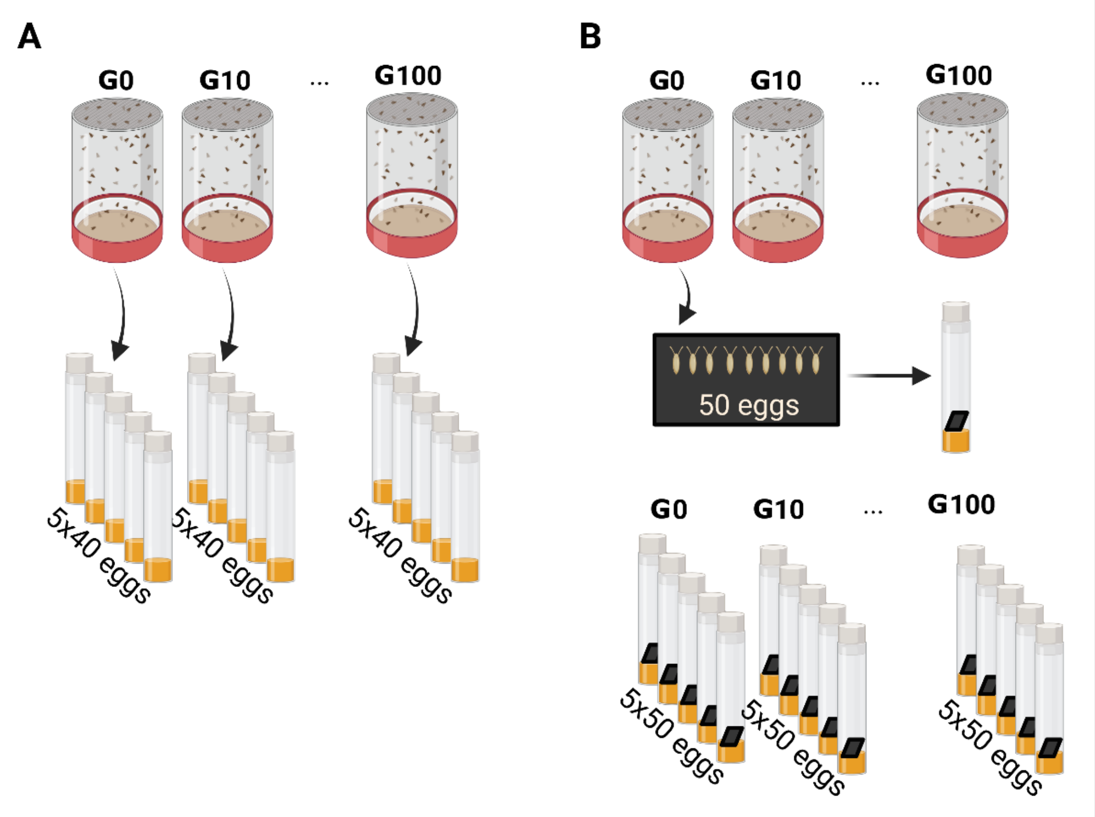

***Figure S7: Experimental design for the viability and hatchability assays.***

*A) For the viability experiment, adults were allowed to lay eggs for a few hours in the morning, and the collected progeny were distributed into five replicate vials with 40 eggs for each condition. The number of emerging adults per vial was recorded to estimate viability. B) For the hatchability experiment, 50 eggs were collected from each cage, transferred into replicate vials, and monitored for hatching success by observation of the eggs laid on a black piece of paper. Hatchability was estimated as the proportion of eggs that successfully hatched across replicates.*

| Population | Sex | Temperature | Effective number |
|---|---|---|---|
| G0 | F | 18 | 120 |
| | | 22 | 111 |
| | M | 18 | 78 |
| | | 22 | 100 |
| G10 | F | 18 | 130 |
| | | 22 | 103 |
| | M | 18 | 134 |
| | | 22 | 102 |
| G31 | F | 18 | 113 |
| | | 22 | 137 |
| | M | 18 | 111 |
| | | 22 | 93 |
| G73 | F | 18 | 103 |
| | | 22 | 100 |
| | M | 18 | 101 |
| | | 22 | 80 |
| G100 | F | 18 | 115 |
| | | 22 | 115 |
| | M | 18 | 120 |
| | | 22 | 90 |

***Table S15: Number of flies assayed for longevity in the TE-accumulation populations under different temperatures.***

*For each population, the table reports the sex of the flies (F: female, M: male), the rearing temperature (°C), and the effective number of individuals for which longevity data were collected.*

| Population | Sex | Medium | Effective number |
|---|---|---|---|
| G0 | F | Agar | 104 |
| | | -Yeast | 150 |
| | | Complete | 67 |
| | | -Sugar | 52 |
| G0 | M | Agar | 100 |
| | | -Yeast | 108 |
| | | Complete | 62 |
| | | -Sugar | 53 |
| G10 | F | Agar | 105 |
| | | -Yeast | 100 |
| | | Complete | 64 |
| | | -Sugar | 57 |

| | | | |
|---|---|---|---|
| G10 | M | Agar | 117 |
| | | -Yeast | 119 |
| | | Complete | 56 |
| | | -Sugar | 59 |
| G31 | F | Agar | 107 |
| | | -Yeast | 150 |
| | | Complete | 97 |
| | | -Sugar | 62 |
| G31 | M | Agar | 104 |
| | | -Yeast | 120 |
| | | Complete | 51 |
| | | -Sugar | 65 |
| G73 | F | Agar | 102 |
| | | -Yeast | 129 |
| | | Complete | 80 |
| | | -Sugar | 56 |
| G73 | M | Agar | 103 |
| | | -Yeast | 124 |
| | | Complete | 55 |
| | | -Sugar | 51 |
| G100 | F | Agar | 106 |
| | | -Yeast | 121 |
| | | Complete | 82 |
| | | -Sugar | 56 |
| G100 | M | Agar | 117 |
| | | -Yeast | 104 |
| | | Complete | 60 |
| | | -Sugar | 60 |

***Table S16: Number of flies assayed for longevity in the TE-accumulation populations on different rearing media.***

*For each population, the table reports the sex of the flies (F: female, M: male), the rearing media, and the effective number of individuals for which longevity data were collected. The "Complete" medium contains both sugar and yeast. The "-Yeast" medium was identical to the complete medium but without yeast, while the "-Sugar" medium lacked sugar. The "Agar" medium consists only of water and preservative, and thus provided no nutritional value.*

| Population | Sex | Concentration | Effective number |
|---|---|---|---|
| G0 | F | 15mM | 98 |
| | | 25mM | 103 |
| | M | 15mM | 99 |

| | | | |
|---|---|---|---|
| | | 25mM | 105 |
| G10 | F | 15mM | 101 |
| | | 25mM | 106 |
| | M | 15mM | 101 |
| | | 25mM | 100 |
| G31 | F | 15mM | 100 |
| | | 25mM | 102 |
| | M | 15mM | 97 |
| | | 25mM | 106 |
| G73 | F | 15mM | 102 |
| | | 25mM | 103 |
| | M | 15mM | 92 |
| | | 25mM | 101 |
| G100 | F | 15mM | 104 |
| | | 25mM | 103 |
| | M | 15mM | 111 |
| | | 25mM | 106 |

***Table S17: Number of flies assayed for longevity in the TE-accumulation populations exposed to variable concentrations of paraquat.***

*For each population, the table reports the sex of the flies (F: female, M: male), the paraquat concentration (15mM and 25mM), and the effective number of individuals for which longevity data were collected.*

### References

Chakraborty, Mahul, J. J. Emerson, Stuart J. Macdonald, and Anthony D. Long. 2019. “Structural Variants Exhibit Widespread Allelic Heterogeneity and Shape Variation in Complex Traits.” *Nature Communications* 10 (1): 4872. https://doi.org/10.1038/s41467-019-12884-1.

Chakraborty, Mahul, Nicholas W. VanKuren, Roy Zhao, Xinwen Zhang, Shannon Kalsow, and J. J. Emerson. 2018. “Hidden Genetic Variation Shapes the Structure of Functional Elements in Drosophila.” *Nature Genetics* 50 (1): 20–25. https://doi.org/10.1038/s41588-017-0010-y.

Groza, Cristian, Xun Chen, Travis J. Wheeler, Guillaume Bourque, and Clément Goubert. 2024. “A Unified Framework to Analyze Transposable Element Insertion Polymorphisms Using Graph Genomes.” *Nature Communications* 15 (1): 8915. https://doi.org/10.1038/s41467-024-53294-2.

Liu, Yan-Nan, Jian-Jun Gao, Xiao-Lin Zhuang, Dong-Dong Wu, and Yan-Bo Sun. 2025. “Near Complete Assembly of Drosophila Melanogaster Canton S Strain Genome.” *Nature Communications*, ahead of print, December 3. https://doi.org/10.1038/s41467-025-67031-w.

Massouras, Andreas, Sebastian M. Waszak, Monica Albarca-Aguilera, et al. 2012. “Genomic Variation and Its Impact on Gene Expression in Drosophila Melanogaster.” *PLOS Genetics* 8 (11): e1003055. https://doi.org/10.1371/journal.pgen.1003055.

Mohamed, Mourdas, Nguyet Thi-Minh Dang, Yuki Ogyama, et al. 2020. “A Transposon Story: From TE Content to TE Dynamic Invasion of Drosophila Genomes Using the Single-Molecule Sequencing Technology from Oxford Nanopore.” *Cells* 9 (8): 8. https://doi.org/10.3390/cells9081776.
